# Nutrient Limitation Sensitizes *Pseudomonas aeruginosa* to Vancomycin

**DOI:** 10.1101/2023.04.10.536232

**Authors:** Derek C. K. Chan, Katherine Dykema, Mahrukh Fatima, Hanjeong Harvey, Ikram Qaderi, Lori L. Burrows

## Abstract

Traditional antibacterial screens rely on growing bacteria in nutrient-replete conditions which are not representative of the natural environment or sites of infection. Instead, screening in more physiologically relevant conditions may reveal novel activity for existing antibiotics. Here, we screened a panel of antibiotics reported to lack activity against the opportunistic Gram-negative bacterium, *Pseudomonas aeruginosa,* under low-nutrient conditions, and discovered that the glycopeptide vancomycin inhibited growth of *P. aeruginosa* at low micromolar concentrations through its canonical mechanism of action, disruption of peptidoglycan cross-linking. Spontaneous vancomycin-resistant mutants had activating mutations in the sensor kinase of the two-component CpxSR system, which induced cross-resistance to almost all classes of β-lactams, including the siderophore antibiotic cefiderocol. Other mutations that conferred vancomycin resistance mapped to WapR, an α-1,3-rhamnosyltransferase involved in lipopolysaccharide core biosynthesis. A WapR P164T mutant had a modified LPS profile compared to wild type that was accompanied by increased susceptibility to select bacteriophages. We conclude that screening in nutrient-limited conditions can reveal novel activity for existing antibiotics and lead to discovery of new and impactful resistance mechanisms.

**For Table of Contents Use Only:** 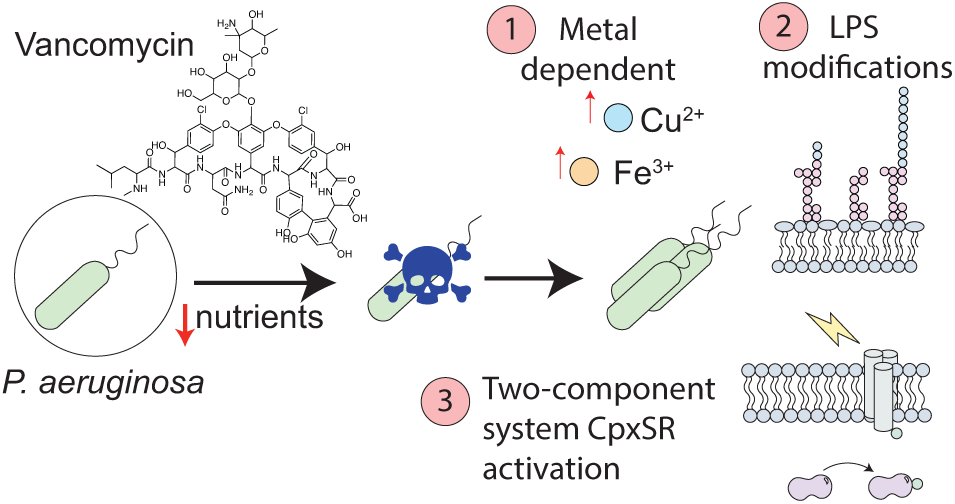

---

The outer membrane (OM) of Gram-negative bacteria provides intrinsic resistance to many antibiotics by reducing uptake. With some exceptions, the OM excludes large and charged molecules with a size-exclusion limit of ∼600 Da^1^. The OM of the Gram-negative opportunistic pathogen *P. aeruginosa* is considered particularly impenetrable, up to 100-fold less permeable compared to that of *Escherichia coli*^2^. This characteristic, coupled with multiple efflux systems which extrude molecules that enter the cell, make *P. aeruginosa* infections difficult to treat. A better understanding of how antibiotics cross the OM may be informative for the development of new strategies to increase drug uptake.

A major contributor to OM impermeability is lipopolysaccharide (LPS), found in the outer leaflet. LPS is composed of a lipid A anchor, an inner and outer core oligosaccharide, and O-antigen of varying length and chemical composition. LPS contributes to membrane stability^3^, motility^4, 5^, biofilm formation^6^_-8_, and antibiotic resistance^9^. LPS is highly negatively charged due to the presence of phosphate groups on lipid A and the inner core region^10^. Divalent cations such as Mg^2+^ and Ca^2+^ are essential to neutralize the negative charge and maintain barrier integrity^11^. Some antibiotics, such as cationic peptides colistin and polymyxin B, can compete with divalent cations for LPS binding, leading to OM permeabilization^11, 12^. The macrolide, azithromycin, acts through a similar mechanism against *P. aeruginosa*^13^. However, bacteria can modify their LPS composition to gain resistance. In *P. aeruginosa,* mutations in the sensor kinase of the two-component PmrAB system leads to activation of multiple genes, including the *arn* operon which catalyzes the covalent addition of 4-amino-4-deoxy-L-arabinose to lipid A^10^. This addition of sugars reduces the binding of cationic antimicrobial peptides. In *E. coli*, mutations in the O-antigen ligase *waaL* lead to incorporation of peptidoglycan precursors that bind vancomycin to prevent entry of the antibiotic into the cell^14^. LPS also serves as a primary receptor for many bacteriophages. In response, bacteria can modify their LPS composition to prevent phage attachment^15, 16^. Phages can counter resistance by encoding enzymes that modify host LPS, to facilitate their own uptake or that of related phages. Overall, LPS composition plays an important role in antibiotic and phage susceptibility.

Once an antibiotic gets inside the cell, bacteria respond to the resulting stress in ways that are not yet fully understood. For example, multiple two-component regulatory systems can be activated in response to specific stimuli^17–20^. Typical two-component systems are composed of a sensor kinase and a response regulator. The sensor can have both kinase (activating) and phosphatase (deactivating) activities that control the phosphorylation state of the response regulator. The response regulator binds upstream of various genes to modulate their expression. Among the best-characterized two-component systems is CpxA-CpxR in *E. coli*, which is activated in response to misfolded proteins in the periplasm^21^, upon overexpression of NlpE^22^, or loss of the L,D-transpeptidase LdtD^23^. *P. aeruginosa* has an orthologous system – CpxSR^24^; however, the exact repertoire of genes regulated by CpxR and the effects downstream of its activation are not yet fully understood. Activation of the Cpx system is involved in antibiotic resistance through the regulation of porin^25^ and efflux pump expression^26^, plus other mechanisms that remain to be discovered.

We previously discovered that the thiopeptide antibiotics, thiostrepton and thiocillin, cross the OM of *P. aeruginosa* using siderophore transporters that are upregulated in low-iron media^27–30^. The thiopeptides synergized with the FDA-approved iron chelator deferasirox (DSX). Prior to our work, the thiopeptides were considered to lack antipseudomonal activity^31, 32^ because susceptibility assays are typically conducted in nutrient-replete media. These findings suggested there may be other large natural product antibiotics that could cross the OM in nutrient-depleted conditions. In this work, we screened existing antibiotics reported to have poor or no activity against *P. aeruginosa* in low-nutrient conditions. We found that the glycopeptide vancomycin had low micromolar activity against *P. aeruginosa* and that its activity was iron-and copper-dependent. Analysis of vancomycin-resistant mutants revealed mutations that activated CpxSR, concomitantly inducing resistance to ý-lactam antibiotics. Further, we identified a mutation in a LPS glycosyltransferase that decreased susceptibility to vancomycin and azithromycin, another large natural product antibiotic, but unexpectedly increased susceptibility to select bacteriophages. These data provide insight on the mechanism of vancomycin uptake in *P. aeruginosa* and describe resistance mechanisms that can promote changes in resistance to other drug classes or therapeutic alternatives, with implications for uptake of antibiotics across the OM barrier and effective treatment of *P. aeruginosa* infections.

## Results

### Vancomycin inhibits *P.* aeruginosa growth in an iron and copper-dependent manner

To identify natural products besides the thiopeptides with possible antipseudomonal activity, we screened 21 commercially available natural product antibiotics at 10μg/mL for activity against *P. aeruginosa* PA14 in nutrient-limited 10:90 (10% lysogeny broth: 90% phosphate-buffered saline) medium in the presence of the iron chelator, deferasirox (64μg/mL) (**Fig. 1A, Supplementary Table S1**). The 21 antibiotics were selected because they were 1) reported to lack antimicrobial activity against *P. aeruginosa* and 2) had a molecular weight >600 kDa, precluding their passive diffusion through porins^33, 34^. Six of the 21 compounds inhibited the growth of *P. aeruginosa* in 10:90 (**Supplementary Table S1**). We focused here on vancomycin because it is used clinically to treat *Staphylococcus aureus* infections that are often associated with *P. aeruginosa* in the cystic fibrosis lung^35–38^. The mechanism of action of vancomycin has also been extensively studied using Gram positive species.

**Figure 1.**
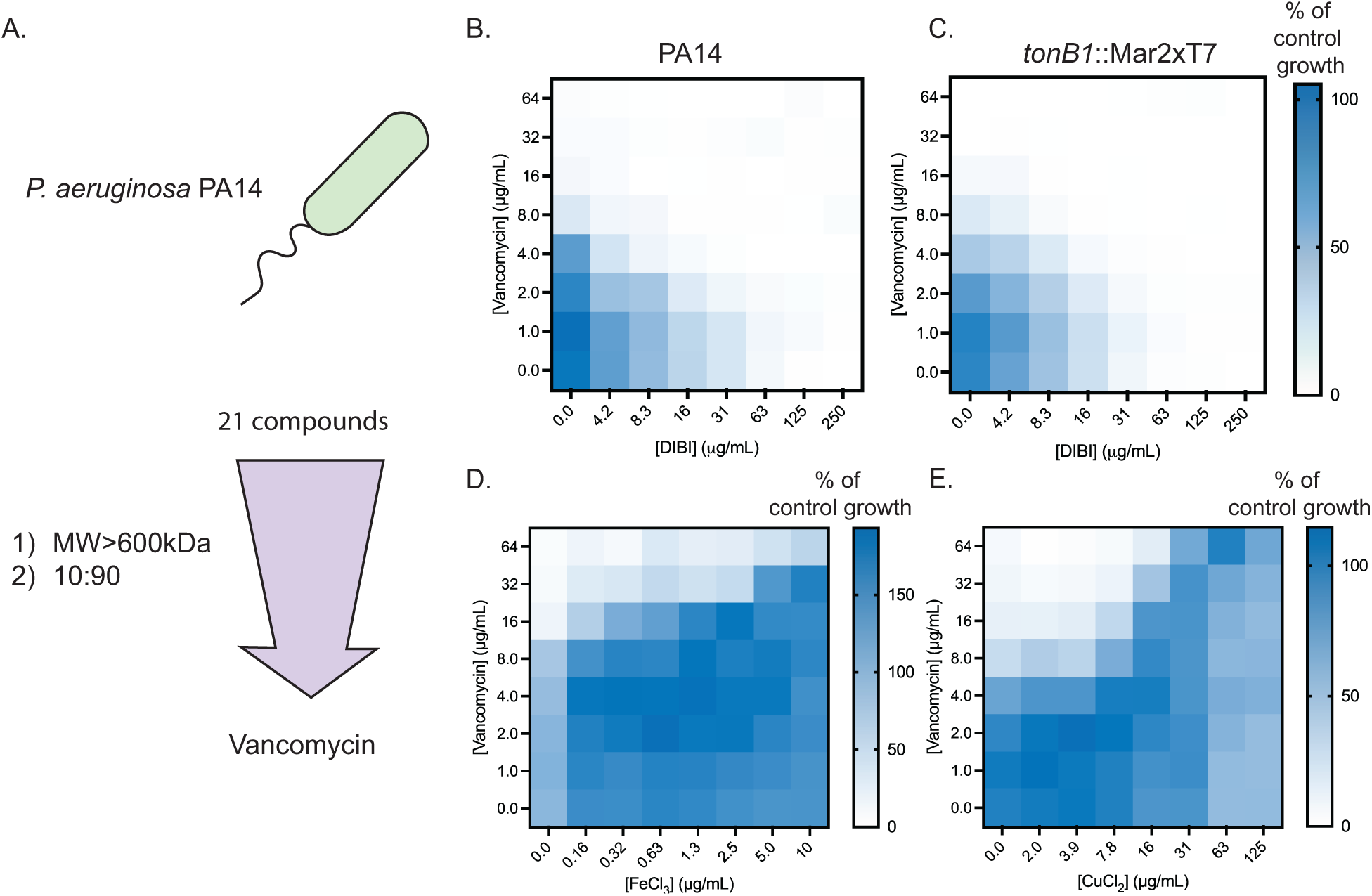
Vancomycin inhibits *P. aeruginosa* under nutrient-limited conditions. **A.** Schematic for screening compounds against *P. aeruginosa*. Checkerboard assays with vancomycin + DIBI against **B.** PA14 and **C.** *tonB1*::Mar2xT7. The darker the shade of blue, the more growth observed. White squares indicate a lack of growth observed. Checkerboard assays with vancomycin and **D.** FeCl_3_ and **E.** CuCl_2_. All checkerboards were averaged from three independent biological replicates.

We first confirmed that vancomycin had activity in combination with other iron chelators using the high-molecular-weight polymeric iron chelator DIBI that is unable to enter cells (∼9,000 kDa)^39^. Similar to the results with DSX, vancomycin synergized with DIBI, further supporting the connection between iron limitation and antibiotic activity (**Fig. 1B**). Vancomycin alone had a minimal inhibitory concentration (MIC) of 16 μg/mL (11 μM) in 10:90, but the addition of 8.3 μg/mL of DIBI reduced the MIC of vancomycin to 4 μg/mL. DIBI alone has a MIC of 63 μg/mL (7 μM) in 10:90. Since the thiopeptides thiostrepton and thiocillin can cross the OM using siderophore transporters, we tested synergy between vancomycin and DIBI against a *tonB1* transposon mutant. The TonB1-ExbBD complex energizes uptake of ligands through OM siderophore transporters and *tonB1* mutants are thiopeptide-resistant^29^. Susceptibility of the *tonB1* mutant to the combination of vancomycin + DIBI was similar to that of the wild type (WT), suggesting that vancomycin uptake did not rely on TonB1-dependent transporters or that loss of *tonB1* was insufficient to reduce activity (**Fig. 1C**). Since vancomycin susceptibility was iron-dependent, we tested the effects of adding back various metal salts, including FeCl_3_, CuCl_2_, MgCl_2_, and CaCl_2_ into 10:90. We predicted that FeCl_3_ would antagonize vancomycin susceptibility because of the observed sensitization by iron chelators. Further, we predicted that MgCl_2_ and CaCl_2_ may have a similar effect because the divalent cations stabilize the highly negative charge of the LPS and thus OM integrity. As expected, iron supplementation antagonized vancomycin susceptibility (**Fig. 1D**). Interestingly, copper supplementation also resulted in antagonism (**Fig. 1E**), but neither MgCl_2_ and CaCl_2_ antagonized vancomycin susceptiblity to the same extent (**Supplementary Fig. S1**). These results suggest that vancomycin susceptibility is mainly Fe^3+^ and Cu^2+^ dependent and that the antibiotic was not simply entering the cell due to compromised OM integrity in 10:90.

### Activity is vancomycin-specific and not due to decreased outer membrane integrity

We verified that the susceptibility to vancomycin was not associated with membrane permeabilization (**Fig. 2A**). 1-N-phenylnaphtylamide (NPN) is a weakly fluorescent dye that poorly permeates the OM; however, upon permeabilization of the OM, the dye binds to phospholipids and strongly fluoresces. Polymyxin B, which disrupts the OM, increased fluorescence whereas vancomycin or piperacillin had no effect on fluorescence compared to the vehicle control.

**Figure 2.**
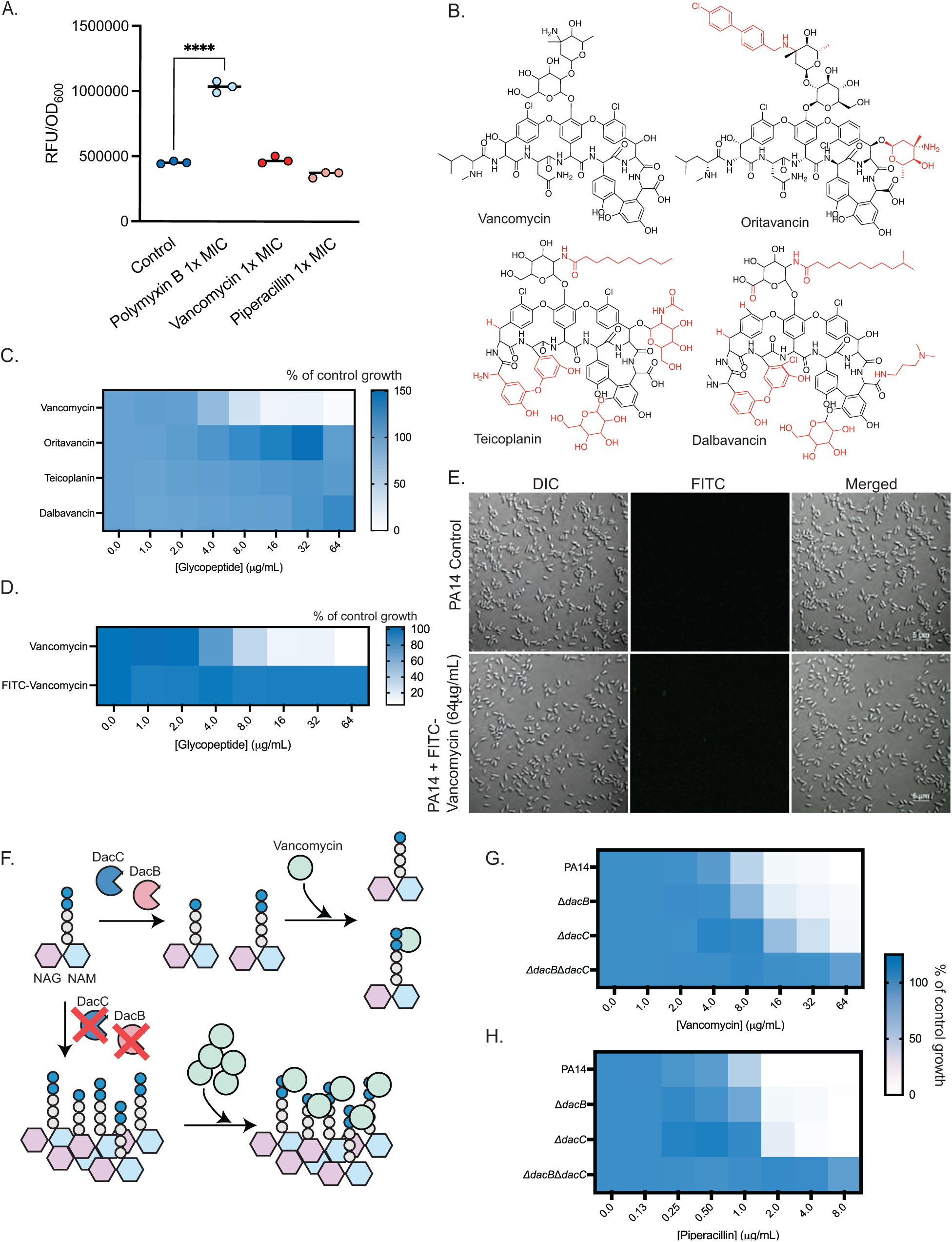
Glycopeptide activity is vancomycin specific and vancomycin acts through its canonical mechanism of action. **A.** NPN assay with PA14 in 10:90 with polymyxin B (1X MIC = 1μg/mL), vancomycin (1X MIC = 16μg/mL), and piperacillin (1X MIC = 8μg/mL). NPN fluorescence was measured with excitation at 350nm and emission at 420nm. Values for individual biological replicates are shown. Statistical analysis was calculated using a one-way ANOVA followed by Dunnett’s multiple comparison test. ****, p<0.0001. **B.** Structures of vancomycin, oritavancin, teicoplanin, and dalbavancin. Structural compared to vancomycin are highlighted in red. **C.** MIC assay with vancomycin and the three glycopeptides against PA14 in 10:90. **D.** MIC assay with FITC-vancomycin against PA14 in 10:90. **E.** Representative fluorescent microscopy images of cells treated with FITC-vancomycin. Scale bar = 5μm. **F.** Schematic for vancomycin (green circle) binding to the D-Ala-D-Ala moiety of the pentapeptide of lipid II. DacB and DacC remove the terminal D-Ala from the pentapeptide stem to form the tetrapeptide. The cell has both penta-and tetrapeptides and vancomycin can bind to the pentapeptides through H-bonding with the D-Ala-D-Ala residues. In the absence of DacB and DacC, the pool of pentapeptides increases, thus more vancomycin is required to inhibit transpeptidation. MIC assays of PA14, Δ*dacB,* Δ*dacC,* and Δ*dacB*Δ*dacC* treated with **G.** vancomycin and **H.** piperacillin. All MIC assays were conducted in 10:90 and averaged from three independent biological replicates.

We next tested if other glycopeptides could inhibit *P. aeruginosa* growth in 10:90 (**Fig. 2B**). Teicoplanin, oritavancin, and dalbavancin all lacked activity up to 64 μg/mL (**Fig. 2C**). We also tested FITC-vancomycin, a fluorescent derivative, and saw no inhibition (**Fig. 2D**). Using microscopy we saw no fluorescence in cells, indicating that the fluorescent analog could likely not cross the OM (**Fig. 2E**). These results show that the activity is vancomycin-specific and sensitive to structural modifications.

Vancomycin is primarily bacteriostatic against *P. aeruginosa* but bactericidal during early exponential phase (**Supplementary Fig. S2A**). At 1X MIC (16 μg/mL), where no growth was observed, cells grew similarly to the WT until early exponential phase, when there was a decrease in OD_600_, indicative of bactericidal activity. Time-kill curves showed that at 1X MIC, there was a 1 log increase in CFU/mL compared to the starting inoculum after 24 h (**Supplementary Fig. 2B**). At 4X MIC, vancomycin was bactericidal up to 8 h; however, regrowth occurred by 24 h. These results suggest that while vancomycin has activity under low-nutrient conditions, *P. aeruginosa* can develop resistance. Therefore, we investigated whether vancomycin was acting through its canonical mechanism of action, how it was taken up, and how *P. aeruginosa* could become resistant.

### Loss of DacB and DacC or activation of the CpxSR two-component system leads to vancomycin resistance

Vancomycin binds the D-Ala-D-Ala moiety of peptidoglycan pentapeptide stems, resulting in decreased crosslinking and impairment of cell wall integrity^40^. Therefore, we predicted that mutations that increase pentapeptide levels would result in decreased susceptibility to vancomycin. Previous studies reported that loss of the D,D-carboxypeptidases, DacB (penicillin binding protein 5, PBP5) and DacC (PBP4), increased the pool of pentapeptides in *P. aeruginosa* (**Fig. 2F**)^41^. Deletion of *dacB* also upregulates expression of the chromosomally-encoded beta-lactamase, AmpC, increasing the MIC for beta lactams such as piperacillin although the degree of AmpC stimulation depends on the strain^42, 43^. Consistent with these data, MICs for both vancomycin and piperacillin increased in a *dacB dacC* double mutant, suggesting that vancomycin was acting through its expected mechanism of action (**Fig. 2GH**).

To understand potential mechanisms of uptake and resistance to vancomycin in *P. aeruginosa*, we screened a library of non-redundant *P. aeruginosa* Mar2xT7 transposon insertion mutants at 64 μg/mL vancomycin (4X MIC) to identify mutants with increased resistance (**Fig. 3A**). Only three mutants were reproducibly resistant: *PA14_64520*, *PA14_12360*, and *PA14_12370*, each with MICs of 64 μg/mL. *PA14_64520* encodes a bacterioferritin contributing to iron homeostasis. *PA14_12360* and *PA14_12370* are tandem uncharacterized genes, suggesting a potential interaction between their protein products or possible polar effects on *PA14_12370* from the transposon insertion in *PA14_12360*. The relationship between the products of these genes and their relation to vancomycin uptake or mechanisms of resistance was unclear; therefore, we also raised spontaneously vancomycin-resistant mutants in both liquid (up to 64 μg/mL) and solid media formats (128-256 μg/mL) (**Fig. 3B**). We sequenced the vancomycin-resistant mutants arising from liquid cultures, which all mapped to the two-component system sensor CpxS, causing a T163P mutation. Interestingly, all mutants raised on solid media with 256 μg/mL vancomycin also had mutations in *cpxS*, suggesting that the CpxSR system is required for high level vancomycin resistance. In *E. coli,* at least 10 mutations in the CpxS ortholog, CpxA, have been shown to be activating mutations. We aligned the AlphaFold2 structural models of CpxA and CpxS, which showed that CpxS T163P is located at transmembrane helix 2, a position similar to those of 5 CpxA activating mutations (**Fig. 4AB**). All the other mutations identified are also located in positions similar to known CpxA activating mutations (**Fig. 4C**).

**Figure 3.**
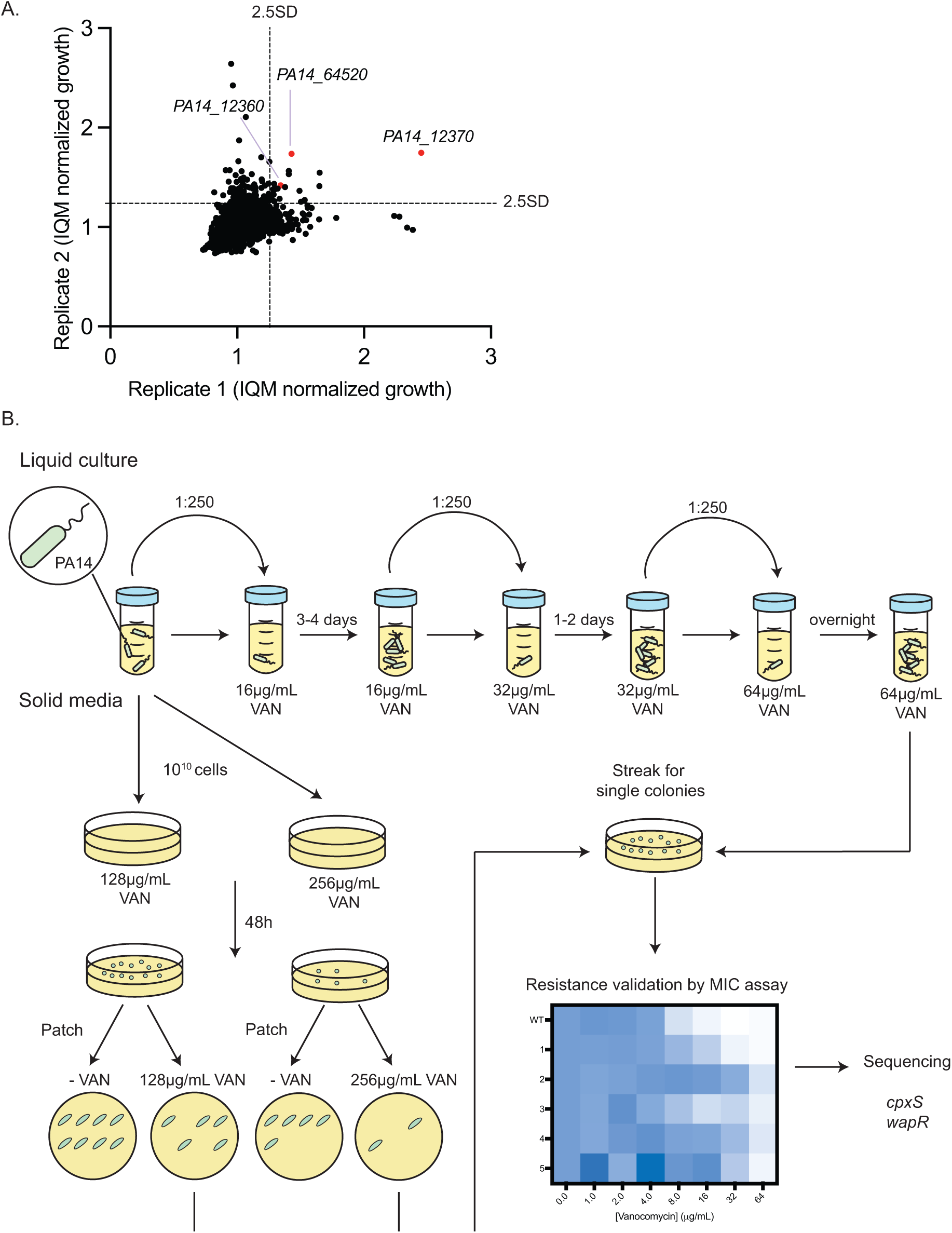
Identifying genes important for vancomycin resistance. **A.** Replica plot of interquartile-mean normalized growth of the PA14 transposon library screened at 64μg/mL vancomycin in 10:90. Mutants with growth >2.5 standard deviations (SD) above the mean were considered as resistant. Validated resistant mutants are highlighted in red and labeled with their gene name. **B.** Schematic for raising spontaneous vancomycin-resistant mutants in liquid and solid media – Vancomycin (VAN).

**Figure 4.**
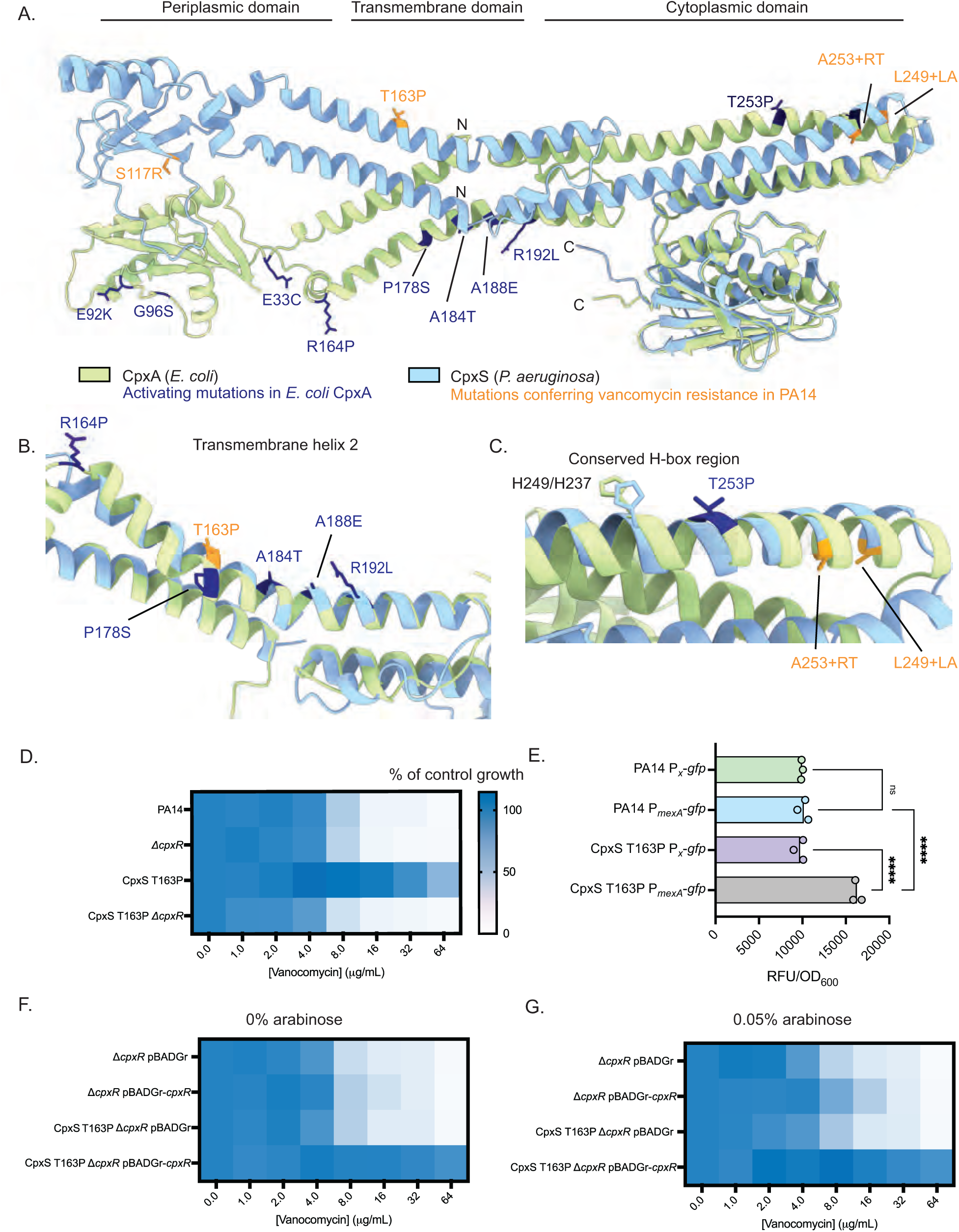
Point mutations in CpxS confer resistance to vancomycin. **A.** AlphaFold2 predictions of *E. coli* CpxA (green) and *P. aeruginosa* CpxS (blue) overlayed. Activating mutations in *E. coli* CpxA are highlighted in dark blue whereas mutations in CpxS in spontaneous vancomycin-resistant mutants are indicated in orange. Structural alignment of **B.** transmembrane helix 2 and **C.** transmembrane helix 1 near the conserved H-box region between CpxA (green) and CpxS (blue). **D.** MIC assay of PA14, Δ*cpxR,* CpxS T163P, and CpxS T163P Δ*cpxR* treated with vancomycin. **E.** GFP-reporter assay of PA14 WT versus CpxS T163P using a promoterless control or P*_mexA_* to drive GFP expression ns: not significant; ****: p<0.0001. Statistics were calculated by one-way ANOVA followed by Dunnett’s multiple comparison test in Prism. Individual values for each biological replicate are shown. Δ*cpxR* and CpxS T163P Δ*cpxR* complemented with empty vector (pBADGr) or a WT copy of *cpxR,* and treated with vancomycin in 10:90 **F.** with no arabinose or **G.** 0.05% arabinose. All assays were conducted in 10:90 and results shown are averaged from three independent biological replicates.

We focused on CpxS T163P because it was recovered from both liquid and solid media cultures. The CpxS T163P mutant was regenerated in a WT background via allelic exchange and we confirmed that it was resistant to vancomycin (**Fig. 4D**) with an MIC > 64μg/mL. We also complemented a *cpxS*::Mar2xT7 mutant with WT PA14 CpxS or CpxS T163P and tested for vancomycin susceptibility (**Supplementary Fig. S3**). The *cpxS* mutant had WT susceptibility to vancomycin with a MIC of 16 μg/mL. Expression of CpxS T163P, but not empty vector or WT CpxS, restored resistance (MIC = 64 μg/mL). In *E. coli*, mutations in the CpxAR system can be activating or repressing; therefore, to learn whether CpxS T163P is an activating mutation, we examined the expression of GFP under the control of the CpxR-responsive *mexA* promoter (**Fig. 4E**). MexA is the periplasmic component of the MexAB-OprM efflux pump, and a previous study showed that CpxR binds upstream of *mexA* at a conserved CpxR box binding motif^26^. We compared the expression of GFP in WT PA14 and the CpxS T163P mutant using either a promoterless reporter or a reporter under control of P*_mexA_*. GFP expression was significantly higher in CpxS T163P compared to the WT or a promoterless control, suggesting that CpxS T163P is an activating mutation.

To confirm that the effect of the CpxS T163P mutation was mediated via the response regulator CpxR, we deleted *cpxR* from WT and CpxS T163P and tested susceptibility to vancomycin (**Fig. 4D**). Δ*cpxR* had WT susceptibility to vancomycin. Further, loss of Δ*cpxR* in the CpxS T163P mutant increased susceptibility to vancomycin to WT levels, suggesting that CpxS T163P activates CpxR to confer resistance. We could also complement the Δ*cpxR* mutants with *cpxR in trans* on an arabinose-inducible plasmid with a gentamicin resistance maker, pBADGr, which restored resistance to vancomycin in the CpxS T163P mutant even without the addition of arabinose, suggesting that low level expression from the leaky promoter was sufficient (**Fig. 4F**). Expression of CpxR *in trans* in Δ*cpxR* increased the MIC slightly in the presence of arabinose (0.05%), suggesting that there is also a dosage-dependent effect even without CpxS activation (**Fig. 4G**), consistent with limited kinase-independent activation of response regulators by small-molecule phosphodonors^44, 45^.

Next, we tested whether activation of the CpxSR response was responsible for observed antagonism between vancomycin + copper and vancomycin + iron. We conducted checkerboard assays with CpxS T163P, Δ*cpxR,* and CpxS T163P Δ*cpxR* challenged with vancomycin + CuCl_2_ and vancomycin + FeCl_3_ (**Supplementary Figure S4**). WT, Δ*cpxR,* and CpxS T163P Δ*cpxR* showed similar profiles of antagonism. CpxS T163P was resistant to vancomycin, but had increased sensitivity to copper compared to other strains. These results suggest that copper and iron antagonize vancomycin activity independently of the CpxSR two-component system.

### Activation of CpxS confers resistance to multiple classes of β-lactams

In *E. coli*, CpxA has been implicated in resistance to multiple classes of antibiotics^46–48;^ therefore, we tested whether CpxS T163P conferred resistance to antibiotics besides vancomycin. The penicillins, monobactams, and cephalosporins showed 4-fold or greater increases in MIC for CpxS T163P compared to the WT (**Fig. 5A**). The mutant also had an elevated MIC of 8-fold for the siderophore antibiotic cefiderocol compared to WT. However, there was no change in MIC for the penems (meropenem and imipenem). Similarly, compounds that inhibit early steps of peptidoglycan synthesis (e.g. D-cycloserine and fosfomycin) or classes of antibiotics with non-cell wall targets showed no change or only a 2-fold change in MIC. These results suggest that CpxS T163P primarily impacts the periplasmic steps of peptidoglycan synthesis, where vancomycin and ý-lactams act. Additionally, the mechanism of resistance may be independent of MexAB-OprM, despite its upregulation in the CpxS T163P mutant, because the MICs for other classes of antibiotics that are substrates for efflux pumps were unchanged. CpxS T163P and WT had similar susceptibility to polymyxin B, suggesting that OM integrity was not impacted.

**Figure 5.**
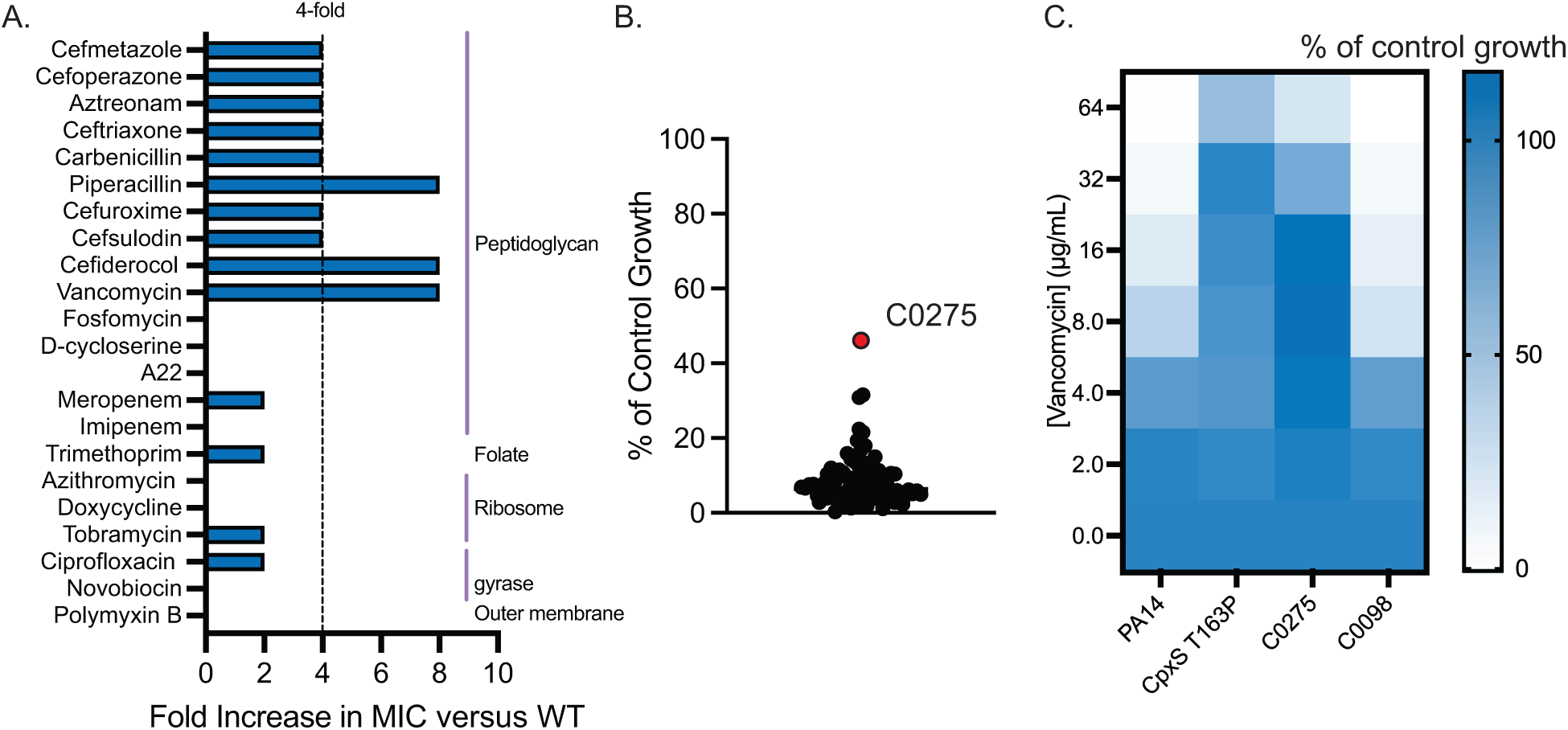
CpxSR activation confers resistance to ý-lactams. **A.** Fold change in MIC of PA14 versus CpxS T163P for the panel of antibiotics tested. Fold changes 4-fold or more are considered resistant. **B.** Screening of the WCC *P. aeruginosa* clinical isolate panel for susceptibility to 64μg/mL vancomycin in 10:90. C0275, a resistant clinical isolate, is highlighted in red. **C.** MIC assay of PA14, CpxS T163P, C0275, and C0098 treated with vancomycin in 10:90. Results for all assays are averaged from three independent biological replicates.

Our data suggested that various CpxS point mutations can activate the CpxSR system. This led us to hypothesize that clinical isolates of *P. aeruginosa* may harbour mutations in CpxS that differ depending on their diversity, prior antibiotic exposure, and the site of infection. We used vancomycin resistance at 64 μg/mL (4x MIC of WT) (**Fig. 5B**) as a filter to uncover potential CpxS-activating mutations in a set of 96 clinical isolates. One isolate C0275 had ∼40% of control growth even at 64 μg/mL vancomycin whereas most isolates had <20% of control growth, which was not observable by eye. C0275 had a MIC for vancomycin >64 μg/mL, similar to CpxS T163P, whereas WT PA14 had a MIC of 16 μg/mL. C0098 was used as a control and had similar susceptibility to WT PA14 (**Fig. 5C**). All the clinical isolates had CpxS sequences similar to those of WT PA14 or PAO1, suggesting that there are other resistance mechanisms that could contribute to the elevated vancomycin MIC in C0275.

### A WapR P164T vancomycin-resistant mutant has altered LPS composition and increased phage sensitivity in nutrient-limited media

We also investigated another spontaneously-resistant mutant raised on 10:90 solid media containing 128 μg/mL vancomycin. This isolate had a mutation in *wapR* leading to a P164T substitution. WapR is an enzyme involved in LPS core oligosaccharide biosynthesis that adds a L-rhamnose (α-1,3-linked) to the outer core to provide an attachment site for O-antigen polymerization. Mutants lacking WapR synthesize rough LPS without O-antigen. To understand the effects of the Pro to Thr mutation, we searched the AlphaFold2 structure of WapR using the Dali server^49, 50^. The best matched identified was a chrondroitin polymerase from *E. coli* K4 (KfoC; PDB: 2Z86)^51^. KfoC is a bifunctional glycosyltransferase that catalyzes the elongation of the chrondroitin chain involved in the synthesis of the extracellular layer of the bacterial capsule. WapR aligned with the A2 domain of K4CP bound to uridine-diphosphate glucuronic acid (UDP-GluUA) (**Fig. 6A**). WapR P164 is located in the region that aligns with the K4CP binding cavity for UDP-GluUA, suggesting that the mutation in WapR may affect its function as an L-rhamnosyltransferase (**Fig. 6B**).

**Figure 6.**
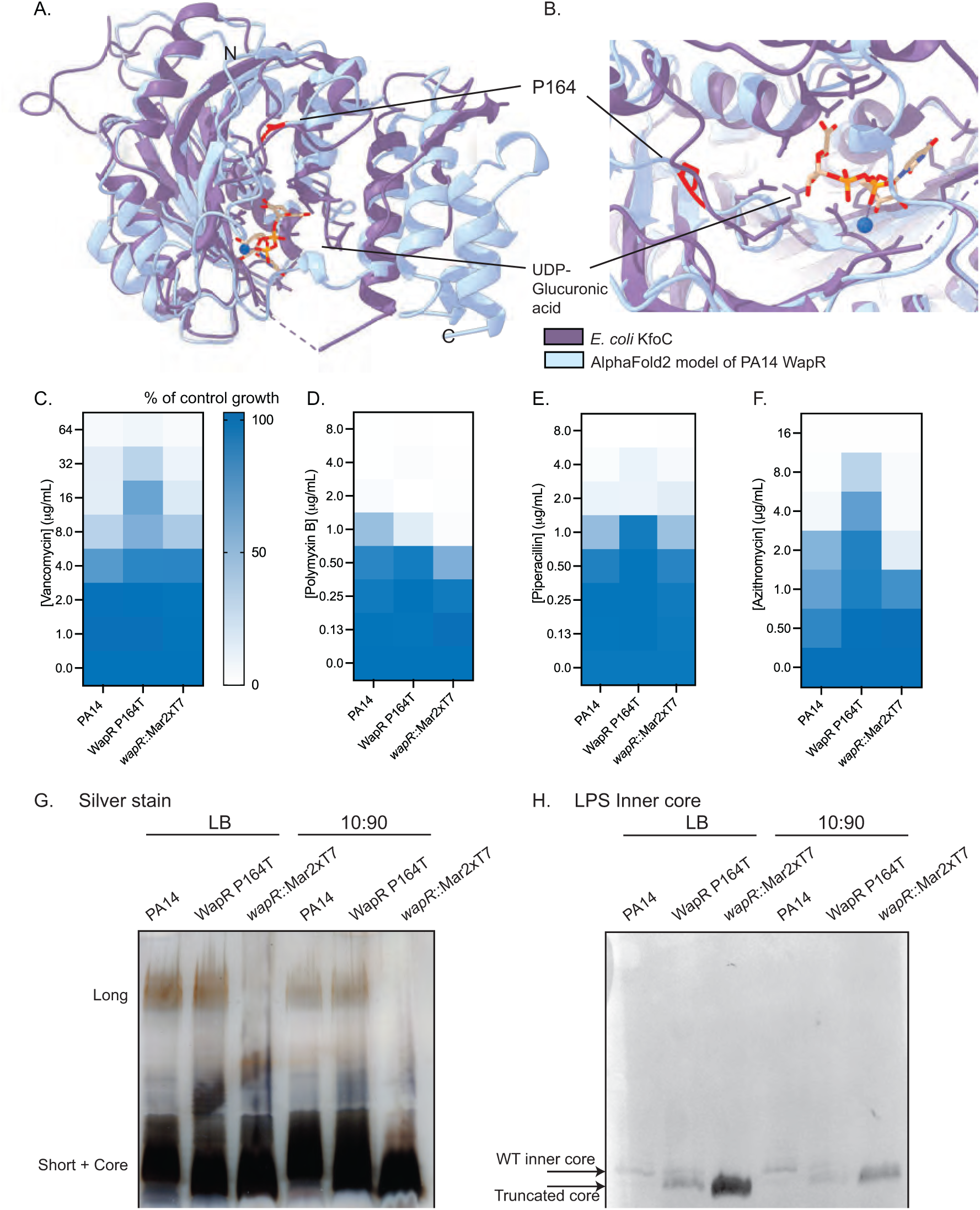
WapR P164T is resistant to vancomycin and azithromycin and has altered LPS profiles. **A.** Overlay of the A2 domain of *E. coli* K4 KfoC (purple) (PDB: 2Z86) and an AlphaFold2 model of PA14 WapR (blue). UDP-glucuronic acid, which binds to KfoC, is highlighted in beige with heteroatoms coloured differently. The blue sphere is Mn^2+^. P164 is highlighted in blue. **B.** A zoomed-in view of the UDP-glucuronic acid binding site of KfoC overlayed with WapR. P164 is highlighted in red. MIC assays of PA14, WapR P164T, and *wapR*::Mar2xT7 treated with **C.** vancomycin, **D.** polymyxin B, **E.** piperacillin, and **F.** azithromycin. Results are averaged from three independent biological replicates. **G.** Silver stain of proteinase K-treated crude lysates for LPS from PA14, WapR P164T, and *wapR*::Mar2xT7. The three strains were grown on LB and 10:90 agar plates. **H.** Western blot of LPS preparations probed with the monoclonal antibody 5c-7-4 (LPS inner core specific)

To test this hypothesis, we tested the susceptibility of WT PA14, WapR P164T, and a *wapR* transposon mutant (*wapR*::Mar2xT7) to vancomycin (**Fig. 6C**). As expected, the WapR P164T mutant had a MIC 4-fold greater than that of the WT. Surprisingly, susceptibility of *wapR*::Mar2xT7 was similar to WT. This result suggests that the WapR P164T mutant may not be a loss-of-function mutation. As a control, we also tested susceptibility of the mutants to polymyxin B (to assess OM integrity) and piperacillin (for peptidoglycan-acting antibiotics) and saw no differences (**Fig. 6DE**). LPS structure modulates antibiotic resistance^10^; therefore, we tested other high-molecular weight antibiotics identified from our initial screen as having activity under nutrient-limited conditions using PA14, WapR P164T, and the *wapR* mutant (**Fig 6F**). Interestingly, the WapR P164T mutant was also 4-fold more resistant to azithromycin compared to the WT, similar to vancomycin. These results suggest that this mutation can also confer resistance to other classes of antibiotics in a LPS-dependent manner.

To further confirm that WapR P164T was not an inactivating mutation, we isolated crude LPS preparations for the three strains grown in LB or 10:90, separated the samples by SDS-PAGE, and visualized LPS by silver staining (**Fig. 6G**). The LPS profile of cells grown in LB were similar to in 10:90 although there appeared to be less long-chain O antigen in 10:90. However in both conditions, WapR P164T makes long-chain length O-antigen that was comparable to the WT while no polymers in that range were present for *wapR*::Mar2xT7. The *wapR*::Mar2xT7 mutant also failed to make short chain length LPS, consistent with previous reports^52^. Additionally, there were differences in the migration pattern of the short + core LPS (**Fig. 6H**). The LPS profiles of the WapR P164T and *wapR*::Mar2xT7 mutants showed that there were bands corresponding to the LPS core region that migrated faster compared to the WT. This observation is consistent with the fact that WapR modifies the LPS core. Therefore, we probed phenol/ethyl ether-extracted LPS preparations with monoclonal antibody 5c-7-4 that recognizes the inner core. For the WT, a single band was present, whereas the *wapR* transposon mutant had a single band of decreased molecular weight, corresponding to a truncated core, consistent with previous observations^53^. However, two bands were seen for the WapR P164T mutant – one higher molecular weight band corresponding to the WT core and a lower molecular weight band corresponding to the *wapR*::Mar2xT7 mutant truncated core. No differences were seen between cells grown in LB or 10:90. Together, these results suggest that the WapR P164T mutation leads to an heterogenous core phenotype, between WT and the *wapR*::Mar2xT7 mutant.

The O antigen is important for antibiotic resistance, but also acts as a receptor for phages to recognize suitable host cells. We tested PA14, WapR P164T, and *wapR*::Mar2xT7 for their susceptibilities to LPS-specific phages P2B9, D6, and E6 (**Fig. 7**). As a control, we also tested three phages (B6, P2A3, and P2F10) that use type IV pili as a receptor, using a Δ*pilA* mutant which is unable to make the major pilin subunit as a negative control (**Fig. 7, Supplementary Figure S5**). We spotted increasing dilutions of each phage on bacterial lawns grown on LB agar and looked for differences in susceptibility. All 6 phages plaqued on WT PA14. When the same set of phages was spotted on WapR P164T, we saw a similar susceptibility pattern to the WT; however, the plaques were clearer and larger. *wapR*::Mar2xT7 was resistant to phages P2B9, D6, and E6, but susceptible to the pilus-specific phages, B6, P2A3, and P2F10, consistent with the requirement for O-antigen binding for infection. The Δ*pilA* mutant was susceptible to phages P2B9, D6, and E6 but resistant to B6, P2A3, and P2F10. The larger plaque sizes for WapR P164T appear to be an LPS-independent phenomenon because the same phenotype was also observed for pilus-dependent phage B6. We also tested the CpxS T163P mutant for phage susceptibility. The titres were similar but the plaques were more turbid compared to WT, suggesting that CpxS activation may impact the efficiency of phage replication.

**Figure 7.**
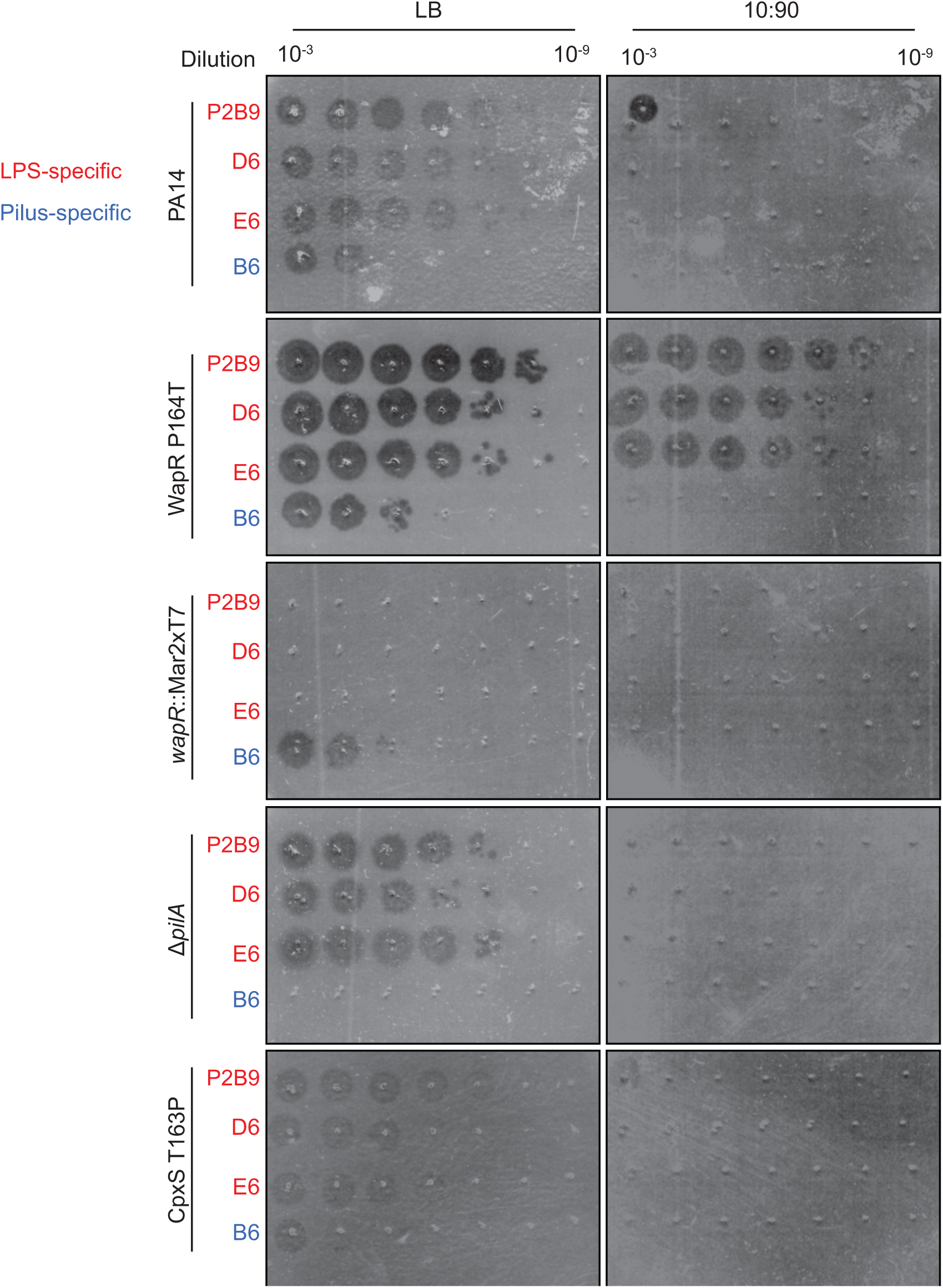
WapR P164T has larger plaques in LB and increased phage susceptibility in 10:90. Phage plaquing assays with PA14, WapR P164T, *wapR*::Mar2xT7, Δ*pilA,* and CpxS T163P in LB and 10:90 agar. Each strain was treated with increasing dilutions of the LPS-specific phages (highlighted in red): P2B9, D6, and E6 and the pilus-specific phage (highlighted in blue) B6. Three biological replicates were conducted and representative plaquing assays are shown.

Despite the differences in plaque sizes between WapR P164T and WT, there were no differences in susceptibility. However, these initial phage assays were conducted in nutrient-replete LB whereas our antibiotic susceptibility assays were done in 10:90. Therefore, we repeated the phage susceptibility assays in 10:90 agar. Interestingly, on that medium PA14 was resistant to LPS-specific phages D6 and E6 and had a 10^5^-fold decrease in susceptibility to P2B9. However, this was not the case for the WapR P164T mutant. We observed similar plaquing and susceptibility patterns of the LPS-specific phages on WapR P164T in LB and 10:90, although it became resistant to pilus-specific phage B6. The *wapR*::Mar2xT7, Δ*pilA,* and CpxS T163P mutants were resistant to all phages tested when grown on 10:90. The differences in phage susceptibility were not due to differences in growth as all strains grew similarly in 10:90 (**Supplementary Fig. S6**). Altogether, these results show that the WapR P164T mutant has altered LPS composition and differences in susceptibility to vancomycin, azithromycin, and phages in rich versus nutrient-limited media.

## Discussion

We found that *P. aeruginosa* is sensitized to the large natural product antibiotic vancomycin when grown in nutrient-limited conditions that may be more representative of the host environment^54^. Vancomycin activity was antagonized by various metals such as iron and copper, but calcium and magnesium – which help to stabilize the negative charge of the OM – were less important for resistance. One potential explanation for the antagonism between vancomycin and copper is that vancomycin can chelate the metal^55, 56^. The functional groups that participate in chelation overlap with those that participate in H-bonding with the D-Ala-D-Ala pentapeptide. Thus, complex formation with copper may inhibit uptake and/or interaction of vancomycin with its target. Copper antagonism of vancomycin activity was independent of the CpxSR pathway, consistent with the chelation hypothesis, although further investigation to support this conclusion is necessary (**Supplementary Fig. S4**). The antagonism of vancomycin activity by iron also suggests that there are alternate mechanisms of resistance yet to be identified but confirms that iron limitation potentiates vancomycin activity.

Our screen of the *P. aeruginosa* PA14 transposon library to identify genes involved in vancomycin resistance yielded only three mutants in poorly-characterized genes. This result suggested that inactivating mutations are not the primary mechanism of *P. aeruginosa* resistance to vancomycin. Instead, isolation of spontaneously-resistant mutants led to the identification of multiple point mutations in CpxS, suggesting that the sensor can be activated through specific substitutions such as T163P, similar to CpxA in *E. coli*^57^. Vancomycin resistance conferred by CpxS T163P was dependent on CpxR, as *cpxR* deletion reduced susceptibility to WT levels.

Detailed examination of the consequences of CpxSR activation will be necessary to understand the transcriptional responses that result in vancomycin resistance. For example, activation of CpxSR may increase the expression of L,D-transpeptidases, implicated in β-lactam resistance because they bypass the effects of D,D-transpeptidase inhibition^58^. In *P. aeruginosa*, 3,3-crosslinking of peptidoglycan catalyzed by L,D-transpeptidases increases in low-nutrient conditions^59^ In *E. coli,* activation of the CpxS homologue CpxA increases expression of L,D-transpeptidase LdtD^23^. The L,D-transpeptidases are not well characterized in *P. aeruginosa,* although there are some candidates (e.g. PA14_27180 and PA14_54810) predicted to have the YkuD-like fold characteristic of this family^60^. In *E. coli*, CpxR also indirectly decreases expression of *dacC* and genes involved in LPS biosynthesis^61^; but whether this changes pentapeptide levels is unclear. Some CpxA-activating mutants have increased abundance whereas others do not^23^. Overall, changes in peptidoglycan composition and remodeling may be one way to confer resistance to both vancomycin and ý-lactams.

Interestingly, activation of CpxSR through the T163P mutation did not increase resistance to other classes of antibiotics. In *E. coli* CpxAR is involved in fosfomycin resistance, by repressing transporters of the antibiotic^46^. These results suggest that the regulatory consequences of activation differ between the two species. *P. aeruginosa* CpxS T163P was identified previously, in a study investigating the efficacy of sequential antibiotic treatment protocols^24^. They found that pre-treating cells with carbenicillin for 15 min increased susceptibility to gentamicin compared to treatment with gentamicin alone. However, among the mutations that conferred resistance to this combination was CpxS T163P, which based on our work may have resulted in increased carbenicillin resistance.

Our finding that CpxS mutations confer resistance to ý-lactams including the recently FDA-approved siderophore-cephalosporin cefiderocol, but not the penems, has significant implications for the treatment of *P. aeruginosa* infections. The target of β-lactams are the PBPs; however, the penem subclass also inhibits the L,D-transpeptidases, which may explain why the CpxS T163P mutant remains susceptible to meropenem and imipenem^58^. Further investigation into the mechanism will be informative. These results, when combined with studies to find additional CpxS-activating mutations, could help guide antibiotic selection for treatment of *P. aeruginosa* infections. For example, in polymicrobial infections, methicillin-resistant *Staphylococcus aureus* (MRSA) is commonly associated with *P. aeruginosa* in chronic wounds and the lungs of cystic fibrosis patients, and vancomycin is used in the treatment of MRSA^54, 62–64^. However, this work shows that vancomycin can also inhibit *P. aeruginosa* under nutrient limiting conditions and select for mutations in CpxS that confer multidrug resistance. Further studies examining the effects of vancomycin on cocultures will be informative to see if this phenomenon could be clinically relevant.

Previous work on *E. coli* showed that cold stress, rather than nutrient limitation, sensitized cells to vancomycin and that truncations in the LPS core restored resistance^65^. They hypothesized that cold stress negatively affected OM integrity, leading to increased susceptibility to vancomycin, while mutants in LPS biosynthesis were better at maintaining the barrier. Resistance was correlated with truncations with the LPS core or production of heterogenous LPS populations. In contrast, our work showed that *wapR*::Mar2xT7, which fails to produce short or long-chain LPS (**Fig. 6G**) and has a truncated core (**Fig. 6H**), had WT levels of vancomycin susceptibility. These results suggest that in *P. aeruginosa,* core truncations alone were insufficient to confer resistance. However, intermediate LPS profiles, as seen with WapR P164T, correlated with resistance. *P. aeruginosa* makes two distinct O-antigens, called A-band or common polysaccharide antigen (CPA) and B-band O-specific antigen (OSA). However, PA14 has a single amino acid substitution G20R in *wbpX* that abolishes CPA production^8^. Therefore, only OSA is produced by the wild type. Another possibility is that the OSA is differentially attached to the core compared to WT LPS. More detailed studies of LPS composition in these strains will help to clarify the mechanism of resistance. Interestingly, WapR P164T was also more resistant to the macrolide antibiotic, azithromycin. Like vancomycin, the activity of azithromycin on *P. aeruginosa* is nutrient dependent and it directly interacts with LPS by displacing divalent cations^13, 66^. However, since the WapR P164T mutant and WT were equally susceptible to polymyxin B, resistance is likely not due to altered OM integrity.

Interestingly, one study reported that a single amino acid mutation in the *E. coli* O-antigen ligase WaaL conferred vancomycin resistance in both WT and mutants with defects in the OM^14^. Resistance was due to modification of LPS with peptidoglycan subunits that bound vancomycin and reduced its uptake^14^. However, this does not appear to be the case with WapR P164T, as we saw no labeling with FITC-vancomycin. As controls, we included methicillin-resistant *S. aureus* MRSA USA 300 which was labeled as expected with FITC-vancomycin (**Supplementary Fig. S7**). WapR P164T was less susceptible to vancomycin and azithromycin, but more susceptible to LPS-specific phages in nutrient-limited 10:90. This combination of phenotypes may be due to altered LPS composition, changes in levels of expression, or exposure of secondary phage receptors. Further investigation into the effects of nutrient limitation on phage replication cycle and burst size will be informative in understanding the mechanism for the observed differences in phage susceptibility. These observations are important considerations for the use of phages as antibiotic alternatives, as many sites of infection can be limiting for key nutrients such as iron^67^.

## Conclusion

In conclusion, we showed that vancomycin – but not other glycopeptides – has antimicrobial activity against *P. aeruginosa* in low-nutrient conditions. Vancomycin-resistant mutants harboured activating mutations in CpxS that also conferred resistance to ý-lactams, including cefiderocol, but not the penems. A mutation in WapR which conferred resistance to vancomycin and azithromycin but increased susceptibility to phages under nutrient-limited conditions was also identified. Our study highlights how screening in low-nutrient conditions can reveal novel activity for existing antibiotics and shows how investigation of resistance mechanisms may help to guide antibiotic therapies for *P. aeruginosa*.

## Experimental Section

### Media and Growth Conditions

All bacteria were cultured overnight in lysogeny broth (LB) at 37°C with 200rpm shaking. Strains with pBADGr and P-*gfp*, cultures were supplemented with 15μg/mL gentamicin. Subcultures (1:500 dilution of overnight cultures) were grown in 10:90 supplemented with or without gentamicin for 3-4hr until at least OD_600_ at 37°C with 200rpm shaking. Arabinose was made as a 20% stock solution in 10:90 and filtered through 0.2μm filters (Fisher Scientific) before diluting into 10:90 for growth assays.

### Bacterial strains, phage, and plasmids

Bacterial strains, phage, and plasmids are listed in **Supplementary Table S2**.

### Molecular Biology

See **Supplementary Table S3** for all primers used in this study. All procedures were conducted as previously described^28, 30^.

### Compounds

**Supplementary Table S4** lists all compounds used in this study. All antibiotic powders were stored at 4°C. Stock solutions were stored at −20°C. Compounds were solubilized in dimethyl sulfoxide or DI H_2_O for assays.

### MIC and Checkerboards assays

Broth microdilution MIC assays and checkerboards were conducted as previously described^27–30^.

### NPN assay

The NPN assay was conducted as previously described with modifications^68^. Briefly, overnight cultures of PA14 in LB were subcultured (1:500 dilution) into 50mL of 10:90 and incubated for 3hrs at 37°C with shaking (200 rpm). Cells were harvested by centrifugation at 3,000G for 5min and washed three times with PBS. Cells were resuspended in 10:90 to a final OD_600_ of 0.1 with 5 μM of carbonyl cyanide m-chlorophenyl hydrazone and 15 μM of NPN. Cells were aliquoted into 96-well black plates with flat clear bottoms (Corning). Antibiotics or vehicle controls were added at 75x the final concentration and fluorescence was read immediately on a BioTek Neo plate reader at excitation and emission wavelengths of 350 nm and 420 nm respectively.

### Fluorescence microscopy

Microscopy was conducted as previously described^28, 30^. Briefly, overnight cultures of PA14 were subcultured in fresh 10:90 (1:500 dilution) and grown at 37°C with shaking (200RPM) for 3-4 hours. Cells were harvested by centrifugation and washed 3X with 1X PBS then incubated with 10μM FITC-vancomycin for 30min at 37°C with shaking (200 rpm). Cells were harvested by centrifugation and washed 3X with 1X PBS before spotting on a 1% agarose pad in 10:90 and imaged with a Nikon A1 confocal microscope through a Plan Apo 60x (NA=1.40) oil objective and acquired with Nikon NIS Elements Advanced Research (V. 5.11.01 64-bit). MRSA USA 300 was grown, labeled, and imaged under the same conditions.

### Transposon library screening and WCC vancomycin susceptibility testing

All mutants from the non-redundant PA14 transposon insertion library^69^ were transferred into 96-well plates containing 150 μL/well liquid LB media and incubated for 16 h at 37⁰C with shaking at 200 rpm. Each mutant was then diluted 1:200 into 10:90 deferrated using FEC-1^70^ to keep the iron concentration consistent between batches of media and incubated for a further 4 h. Mutants were then diluted 1:200 into deferrated 10:90 LB containing 64 μg/mL vancomycin (two replicates) and media without vancomycin (one replicate). Plates were incubated for 16 h at 37⁰C with shaking at 200 rpm, then OD_600_ for each well was recorded. OD_600_ values were normalized to the interquartile mean of each plate and each well position^71^. Growth of >2.5 SD above the mean in the presence of vancomycin was considered a hit. The *P. aeruginosa* WCC was tested in a similar manner except untreated 10:90 was used.

### Spontaneous vancomycin-resistant P. aeruginosa

Spontaneously resistant mutants were selected in liquid and solid media. For liquid media, a PA14 overnight culture was diluted 1:500 into 5 mL cultures of 10:90 LB media containing 16 μg/mL vancomycin, or equivalent volumes of sterile H_2_O. The cultures were incubated at 37⁰C with 200 rpm shaking and inspected daily. When turbidity (growth) was observed, the culture was diluted 1:250 into fresh media containing 2x concentration of compounds. The DIBI concentration was held at 16 μg/mL as higher concentrations prevented growth. Cultures were passaged until growth was observed at 64 μg/mL vancomycin then streaked on LB agar. Single colonies were selected and their genomic DNA sequenced.

For solid media, 10^7^ cells of PA14 subcultured in 10:90 were plated onto 10:90 + 1.5% agar containing 128 and 256 μg/mL in triplicates. Plates were incubated at 37°C for 48h where colonies appeared. Sixteen colonies in total from the 128 and 256 μg/mL plates were patched onto fresh 10:90 plates with and without vancomycin. For example, a colony that grew from a plate containing 128 μg/mL vancomycin would be patched onto a plate without vancomycin and a plate containing 128 μg/mL vancomycin. Mutants that grew on both plates were streaked for single colonies for MIC testing to confirm resistance to vancomycin (>= 4-fold MIC). This process was repeated for the Δ*cpxR* mutant.

### Genomic DNA isolation and sequencing

Genomic DNA of vancomycin-resistant mutants was isolated using Promega Wizard Genomic Isolation Kit. Samples were sent to SeqCenter (Pittsburgh, USA) for Illumina sequencing. FASTA files were processed using FASTQ Groomer, Trimmomatic, FASTQ-interlacer, and FASTQ de-interlacer. Mutations were identified using breseq by comparing to the reference genome (accession number GCF_000404265.1)^72^.

### LPS isolation (whole cell and phenol/ethyl ether extraction)

LPS from PA14, WapR P164T, and *wapR*::Mar2xT7 were isolated as previously described with some modifications^73^. Cells from overnight cultures in LB or 10:90 1.5% agar plates were collected and resuspended in PBS. The OD_600_ was standardized to 2.0 and cells collected again by centrifugation. Cells were resuspended in 150 μL of lysing buffer (2% SDS, 4% 2-mercaptoethanol, 10% glycerol, 0.1M Tris-HCl, pH 6.8) and boiled for 10 min. After cooling to room temperature, proteinase K (NEB) (10 μl of 20mg/mL) was added and incubated at 60°C for 1 h. Proteinase K-treated preparations were used for silver staining. For LPS core samples detected by Western blot, prewarmed phenol solution (60°C) containing 90% phenol (Sigma), 0.1% 2-mercaptoethanol, and 0.2% 8-hydroxyquinoline (Sigma) was added to each lysate (1:1 v/v phenol:lysate) and incubated for 15 min at 60°C. The tubes were then incubated on ice for 10 min before centrifugation (21,000 x g for 5 min). The top aqueous layer was transferred to a fresh tube and 500 μL of ethyl ether solution containing 20 mM Tris-HCl, 1 mM EDTA, pH 8.0 was added. Tubes were centrifuged for 1 min at 21,000 x g and the top ethyl ether layer was removed by aspiration. An equal volume of 2X SDS-PAGE loading buffer was added. All LPS samples were stored at −20°C.

### SDS-PAGE and Western blot

SDS-PAGE and Western blot were conducted as previously described^28, 30^ with some modifications. For the detection of inner LPS core, isolated LPS was separated on a 15% SDS-PAGE gel at 90V for 10 min followed by 200V for 1h. After transferring the LPS to a nitrocellulose membrane and blocking for 1h with 5% skim milk, the blot was incubated overnight with mAb 5c-7-4 (specific for the inner core; 1:100 dilution in PBS)^74^. Isolated LPS was also separated on a 12.5% SDS-PAGE gel at 120V for 1.5hr with the same transferring and blocking steps. The next morning, the blot was washed 3x with PBS for 5 min/wash and incubated with 1:500 α-mouse-alkaline phosphatase in PBS for 1 h. The blot was washed 3x with PBS for 5 min/wash and rinsed briefly with DI H_2_O before detection with BCIP and NBT. Bands appeared after 5-15 mins and imaged with an Azure 400 imaging system.

### Silver stain

Silver stain was conducted as previously described with modifications^73^. Four μL of each LPS preparation were loaded onto a 12.5% polyacrylamide gel and separated for 1.5hr at 120V. The gel was then incubated in EtOH:acetic acid (40%:10%) overnight on a shaking platform at room temperature. The next morning cells were treated with periodic acid solution (40% ethanol: 10% acetic acid: 0.7% periodic acid) for 30 min then washed 3x with DI H_2_O for 10 min each. Silver stain solution (2mL NH_4_OH, 28mL 0.1M NaOH, 115mL DI H_2_O) was added and the gel incubated for 30 min. The gel was then washed two times (6 min/wash). Overwashing led to destaining of the high molecular weight LPS. Gels were developed immediately with developing solution (100 μL 37% formaldehyde, 10 mg citric acid, 200 mL DI H_2_O) until bands could be visualized. Development was stopped by putting the gel in 10% acetic acid. Gels were imaged using an Azure 400 imaging system.

### Phage isolation, purification, and plaquing assays

Environmental phage lysates isolated and amplified using *P. aeruginosa* PA14 were serially diluted (10^-1^ to 10^-8^) in phage buffer (68 mM NaCl, 10 mM Tris-HCl (7.5), 10 mM MgSO_4_, 10 mM CaCl_2_). Five microlitres of each dilution was spotted onto the prepared plates. Phage spots were air dried for 10 min with the lid on. Plates were incubated inverted for 18 h at 37°C. Phage lysate dilutions that produced visible plaques were used for phage plaque purification. Ten microlitres of serially diluted phage lysate and 100 μL of culture were added to 10 ml of top agar (0.6% agar) and poured onto pre-set 1% LB-agar. Plates were incubated inverted for 18 h at 37°C. Plates that had 2-30 plaques were used for plaque purification. A pipette was used to touch the centre of a plaque, followed by resuspension in 100 μL of phage buffer. This process was repeated 2-3 times or until uniform plaque morphology was observed.

Phage plaque assays were conducted as previously described^75^. Briefly, bacteria were grown at 37°C overnight then subcultured in LB or 10:90 (1:100 dilution) and cultured until the OD_600_ reached 0.3 for LB and 0.2 for 10:90. One hundred μL of the LB subculture or 200 μL of the 10:90 culture was mixed with 10 mL of LB + 0.6% agar or 10:90 + 0.6% agar. Phage stocks were serially diluted 10-fold in phage buffer (68 mM NaCl, 10 mM Tris-HCl (7.5), 10 mM MgSO_4_, 10 mM CaCl_2_) and 5 μL of each dilution was spotted. Plates were allowed to dry with the lid on for 10 min, inverted, and incubated at 37°C overnight. The next day plates were imaged. Each experiment was repeated at least three times, representative plates are shown.

### Structural comparisons and phylogenetic analyses

High confidence structural models of CpxA, CpxS, and WapR were generated using AlphaFold2^50^. The AlphaFold2 model of WapR was used to look for similar proteins using Dali^49^. The structure of KfoC was retrieved from the Protein Data Bank (PDB: 2Z86)^51^. Structural alignments were conducted using ChimeraX^76^.

## Supporting Information

Figure S1. Checkerboard assays with PA14 treated with vancomycin and MgCl_2_ or CaCl_2_ Figure S2. Growth curve and time-kill assays of PA14 treated with vancomycin in 10:90

Figure S3. PA14 *cpxS*::Mar2xT7 expressing CpxS T163P in 10:90 + 0.25% arabinose treated with vancomycin

Figure S4. Checkerboard assays with Δ*cpxR*, CpxS T163P, and CpxS T163P Δ*cpxR* with vancomycin and CuCl_2_ or FeCl_3_

Figure S5. WapR P164T treated with pilus-specific phages in LB and 10:90

Figure S6. Growth curves of PA14, WapR P164T, *wapR*::Mar2xT7, Δ*pilA*, and CpxS T163P in 10:90

Figure S7. Fluorescence microscopy images of WapR P164T and MRSA USA 300 treated with FITC-vancomycin

Table S1. Antibiotics screened under low-nutrient conditions against *P. aeruginosa*

Table S2. Strain list Table S3. Primer list

Table S4. Compound source list

## Corresponding Author

**Lori L. Burrows** – *McMaster University, Department of Biochemistry and Biomedical Sciences, Institute for Infectious Disease Research, David Braley Centre for Antibiotic Discovery, L8S 4L8 Hamilton, ON, Canada; https://orcid.org/0000-0003-0838-5040;*

## Authors

**Derek C. K. Chan** – *McMaster University, Department of Biochemistry and Biomedical Sciences, Institute for Infectious Disease Research, David Braley Centre for Antibiotic Discovery, L8S 4L8 Hamilton, ON, Canada; https://orcid.org/0000-0003-0388-5417*

**Katherine Dykema –** *McMaster University, Department of Biochemistry and Biomedical Sciences, Institute for Infectious Disease Research, David Braley Centre for Antibiotic Discovery, L8S 4L8 Hamilton, ON, Canada*

**Mahrukh Fatima** – *McMaster University, Department of Biochemistry and Biomedical Sciences, Institute for Infectious Disease Research, David Braley Centre for Antibiotic Discovery, L8S 4L8 Hamilton, ON, Canada*

**Hanjeong Harvey** – *McMaster University, Department of Biochemistry and Biomedical Sciences, Institute for Infectious Disease Research, David Braley Centre for Antibiotic Discovery, L8S 4L8 Hamilton, ON, Canada*

**Ikram Qaderi** – *McMaster University, Department of Biochemistry and Biomedical Sciences, Institute for Infectious Disease Research, David Braley Centre for Antibiotic Discovery, L8S 4L8 Hamilton, ON, Canada*

## Author Contributions

D.C.K.C. and L.L.B. designed the experiments and wrote the draft. D.C.K.C. performed the fluorescence microscopy, silver stain, western blot, GFP-promoter assays, phage assay, clinical isolate screening and analysis experiments. K.D. and D.C.K.C. conducted the transposon mutant screen. D.C.K.C. designed primers and D.C.K.C. and K.D. made the mutants. D.C.K.C. and K.D. conducted the checkerboards, antibiotic susceptibility assays, and raised spontaneous resistant mutants. M.F. assisted with the primary screen for antibiotics with increased activity under low-nutrient conditions. H.H. made the *pilA* mutant, isolated phages from environmental samples, and assisted with phage plaquing and silver stain assays. I.Q. purified phage P2B9 for phage studies.

## Notes

The authors declare no competing financial interests.

## Acknowledgements

We thank Dr. Bruce Holbein (Fe Pharmaceuticals) for providing DIBI and FEC-1 and Dr. Joseph Lam for anti-LPS core mAb 5c-7-4. This work was supported by a Natural Sciences and Engineering Research Council (NSERC) Discovery Grant RGPIN-2021-04237 to LLB. LLB holds a Tier 1 Canada Research Chair in Microbe-Surface Interactions. DCKC holds a Canadian Institute of Health Research (CIHR) Canada Graduate Scholarship – Doctoral program (CGS-D), and IQ holds a CIHR Masters award (CGS-M). KD held a NSERC Undergraduate Student Research Award.

## Abbreviations Used

(NPN): 1-N-phenylnaphtylamide
(CPA): Common Polysaccharide Antigen
(LPS): Lipopolysaccharide
(MRSA): Methicillin-resistant *Staphylococcus aureus*
(MIC): Minimal inhibitory concentration
(OSA): O-Specific Antigen
(OM): Outer membrane
(PBP): Penicillin Binding Protein
(UDP-GluUA): Uridine-diphosphate glucuronic acid
(WT): Wild type

## References

1. van den Berg, B.; Prathyusha Bhamidimarri, S.; Dahyabhai Prajapati, J.; Kleinekathöfer, U.; Winterhalter, M. Outer-Membrane Translocation of Bulky Small Molecules by Passive Diffusion. Proc. Natl. Acad. Sci. 2015, 112 (23), E2991–E2999. https://doi.org/10.1073/pnas.1424835112.

2. Yoshimura, F.; Nikaido, H. Permeability of *Pseudomonas aeruginosa* Outer Membrane to Hydrophilic Solutes. J. Bacteriol. 1982, 152 (2), 636–642.

3. Kucharska, I.; Liang, B.; Ursini, N.; Tamm, L. K. Molecular Interactions of Lipopolysaccharide with an Outer Membrane Protein from *Pseudomonas aeruginosa* Probed by Solution NMR. Biochemistry 2016, 55 (36), 5061–5072. https://doi.org/10.1021/acs.biochem.6b00630.

4. Toguchi, A.; Siano, M.; Burkart, M.; Harshey, R. M. Genetics of Swarming Motility in *Salmonella enterica* Serovar Typhimurium: Critical Role for Lipopolysaccharide. J. Bacteriol. 2000, 182 (22), 6308–6321.

5. Bowden, M. G.; Kaplan, H. B. The *Myxococcus xanthus* Lipopolysaccharide O-Antigen Is Required for Social Motility and Multicellular Development. Mol. Microbiol. 1998, 30 (2), 275–284. https://doi.org/10.1046/j.1365-2958.1998.01060.x.

6. Penterman, J.; Nguyen, D.; Anderson, E.; Staudinger, B. J.; Greenberg, E. P.; Lam, J. S.; Singh, P. K. Rapid Evolution of Culture-Impaired Bacteria during Adaptation to Biofilm Growth. Cell Rep. 2014, 6 (2), 293–300. https://doi.org/10.1016/j.celrep.2013.12.019.

7. Murphy, K.; Park, A. J.; Hao, Y.; Brewer, D.; Lam, J. S.; Khursigara, C. M. Influence of O Polysaccharides on Biofilm Development and Outer Membrane Vesicle Biogenesis in *Pseudomonas aeruginosa* PAO1. J. Bacteriol. 2014, 196 (7), 1306–1317. https://doi.org/10.1128/JB.01463-13.

8. Hao, Y.; Murphy, K.; Lo, R. Y.; Khursigara, C. M.; Lam, J. S. Single-Nucleotide Polymorphisms Found in the MigA and WbpX Glycosyltransferase Genes Account for the Intrinsic Lipopolysaccharide Defects Exhibited by *Pseudomonas aeruginosa* PA14. J. Bacteriol. 2015, 197 (17), 2780–2791. https://doi.org/10.1128/JB.00337-15.

9. Wang, J.; Ma, W.; Fang, Y.; Liang, H.; Yang, H.; Wang, Y.; Dong, X.; Zhan, Y.; Wang, X. Core Oligosaccharide Portion of Lipopolysaccharide Plays Important Roles in Multiple Antibiotic Resistance in *Escherichia coli*. Antimicrob. Agents Chemother. 65 (10), e00341–21. https://doi.org/10.1128/AAC.00341-21.

10. Chen, H. D.; Groisman, E. A. The Biology of the PmrA/PmrB Two-Component System: The Major Regulator of Lipopolysaccharide Modifications. Annu. Rev. Microbiol. 2013, 67 (1), 83–112. https://doi.org/10.1146/annurev-micro-092412-155751.

11. Sun, J.; Rutherford, S. T.; Silhavy, T. J.; Huang, K. C. Physical Properties of the Bacterial Outer Membrane. Nat. Rev. Microbiol. 2022, 20 (4), 236–248. https://doi.org/10.1038/s41579-021-00638-0.

12. Manioglu, S.; Modaresi, S. M.; Ritzmann, N.; Thoma, J.; Overall, S. A.; Harms, A.; Upert, G.; Luther, A.; Barnes, A. B.; Obrecht, D.; Müller, D. J.; Hiller, S. Antibiotic Polymyxin Arranges Lipopolysaccharide into Crystalline Structures to Solidify the Bacterial Membrane. Nat. Commun. 2022, 13 (1), 6195. https://doi.org/10.1038/s41467-022-33838-0.

13. Imamura, Y.; Higashiyama, Y.; Tomono, K.; Izumikawa, K.; Yanagihara, K.; Ohno, H.; Miyazaki, Y.; Hirakata, Y.; Mizuta, Y.; Kadota, J.; Iglewski, B. H.; Kohno, S. Azithromycin Exhibits Bactericidal Effects on *Pseudomonas aeruginosa* through Interaction with the Outer Membrane. Antimicrob. Agents Chemother. 2005, 49 (4), 1377–1380. https://doi.org/10.1128/AAC.49.4.1377-1380.2005.

14. Grabowicz, M.; Andres, D.; Lebar, M. D.; Malojčić, G.; Kahne, D.; Silhavy, T. J. A Mutant *Escherichia coli* That Attaches Peptidoglycan to Lipopolysaccharide and Displays Cell Wall on Its Surface. eLife 2014, 3, e05334. https://doi.org/10.7554/eLife.05334.

15. Lam, J. S.; Taylor, V. L.; Islam, S. T.; Hao, Y.; Kocíncová, D. Genetic and Functional Diversity of *Pseudomonas aeruginosa* Lipopolysaccharide. Front. Microbiol. 2011, 2, 118. https://doi.org/10.3389/fmicb.2011.00118.

16. Huszczynski, S. M.; Lam, J. S.; Khursigara, C. M. The Role of *Pseudomonas aeruginosa* Lipopolysaccharide in Bacterial Pathogenesis and Physiology. Pathogens 2019, 9 (1), 6. https://doi.org/10.3390/pathogens9010006.

17. Kilmury, S. L. N.; Burrows, L. L. Type IV Pilins Regulate Their Own Expression via Direct Intramembrane Interactions with the Sensor Kinase PilS. Proc. Natl. Acad. Sci. 2016, 113 (21), 6017– 6022. https://doi.org/10.1073/pnas.1512947113.

18. Francis, V. I.; Stevenson, E. C.; Porter, S. L. Two-Component Systems Required for Virulence in *Pseudomonas aeruginosa*. FEMS Microbiol. Lett. 2017, 364 (11), fnx104. https://doi.org/10.1093/femsle/fnx104.

19. Kilmury, S. L. N.; Burrows, L. L. The *Pseudomonas aeruginosa* PilSR Two-Component System Regulates Both Twitching and Swimming Motilities. mBio 2018, 9 (4), e01310–18. https://doi.org/10.1128/mBio.01310-18.

20. Wang, B. X.; Cady, K. C.; Oyarce, G. C.; Ribbeck, K.; Laub, M. T. Two-Component Signaling Systems Regulate Diverse Virulence-Associated Traits in *Pseudomonas aeruginosa*. Appl. Environ. Microbiol. 2021, 87 (11), e03089–20. https://doi.org/10.1128/AEM.03089-20.

21. Isaac, D. D.; Pinkner, J. S.; Hultgren, S. J.; Silhavy, T. J. The Extracytoplasmic Adaptor Protein CpxP Is Degraded with Substrate by DegP. Proc. Natl. Acad. Sci. 2005, 102 (49), 17775–17779. https://doi.org/10.1073/pnas.0508936102.

22. Raivio, T. L.; Leblanc, S. K. D.; Price, N. L. The *Escherichia coli* Cpx Envelope Stress Response Regulates Genes of Diverse Function That Impact Antibiotic Resistance and Membrane Integrity. J. Bacteriol. 2013, 195 (12), 2755–2767. https://doi.org/10.1128/JB.00105-13.

23. Bernal-Cabas, M.; Ayala, J. A.; Raivio, T. L. The Cpx Envelope Stress Response Modifies Peptidoglycan Cross-Linking via the l,d-Transpeptidase LdtD and the Novel Protein YgaU. J. Bacteriol. 2015, 197 (3), 603–614. https://doi.org/10.1128/JB.02449-14.

24. Roemhild, R.; Gokhale, C. S.; Dirksen, P.; Blake, C.; Rosenstiel, P.; Traulsen, A.; Andersson, D. I.; Schulenburg, H. Cellular Hysteresis as a Principle to Maximize the Efficacy of Antibiotic Therapy. Proc. Natl. Acad. Sci. U. S. A. 2018, 115 (39), 9767–9772. https://doi.org/10.1073/pnas.1810004115.

25. Batchelor, E.; Walthers, D.; Kenney, L. J.; Goulian, M. The *Escherichia coli* CpxA-CpxR Envelope Stress Response System Regulates Expression of the Porins OmpF and OmpC. J. Bacteriol. 2005, 187 (16), 5723–5731. https://doi.org/10.1128/JB.187.16.5723-5731.2005.

26. Tian, Z.-X.; Yi, X.-X.; Cho, A.; O’Gara, F.; Wang, Y.-P. CpxR Activates MexAB-OprM Efflux Pump Expression and Enhances Antibiotic Resistance in Both Laboratory and Clinical NalB-Type Isolates of *Pseudomonas aeruginosa*. PLoS Pathog. 2016, 12 (10), e1005932. https://doi.org/10.1371/journal.ppat.1005932.

27. Ranieri, M. R. M.; Chan, D. C. K.; Yaeger, L. N.; Rudolph, M.; Karabelas-Pittman, S.; Abdo, H.; Chee, J.; Harvey, H.; Nguyen, U.; Burrows, L. L. Thiostrepton Hijacks Pyoverdine Receptors To Inhibit Growth of *Pseudomonas aeruginosa*. Antimicrob. Agents Chemother. 2019, 63 (9), e00472–19. https://doi.org/10.1128/AAC.00472-19.

28. Chan, D. C. K.; Burrows, L. L. *Pseudomonas aeruginosa* FpvB Is a High-Affinity Transporter for Xenosiderophores Ferrichrome and Ferrioxamine B. mBio 2022, 0 (0), e03149–22. https://doi.org/10.1128/mbio.03149-22.

29. Chan, D. C. K.; Burrows, L. L. Thiocillin and Micrococcin Exploit the Ferrioxamine Receptor of *Pseudomonas aeruginosa* for Uptake. J. Antimicrob. Chemother. 2021, 76 (8), 2029–2039. https://doi.org/10.1093/jac/dkab124.

30. Chan, D. C. K.; Josts, I.; Koteva, K.; Wright, G. D.; Tidow, H.; Burrows, L. L. A Single Extracellular Loop of FoxA Controls Ligand Specificity, Uptake, and Signaling in Pseudomonas aeruginosa. bioRxiv November 18, 2022, p 2022.11.18.517105. https://doi.org/10.1101/2022.11.18.517105.

31. Su, T. L. Micrococcin. An Antibacterial Substance Formed by a Strain of Micrococcus. Br. J. Exp. Pathol. 1948, 29 (5), 473–481.

32. Kelly, J.; Kutscher, A. H.; Tuoti, F. Thiostrepton, a New Antibiotic: Tube Dilution Sensitivity Studies. Oral Surg. Oral Med. Oral Pathol. 1959, 12 (11), 1334–1339. https://doi.org/10.1016/0030-4220(59)90222-1.

33. Nikaido, H. Molecular Basis of Bacterial Outer Membrane Permeability Revisited. Microbiol. Mol. Biol. Rev. 2003, 67 (4), 593–656. https://doi.org/10.1128/MMBR.67.4.593-656.2003.

34. O’Shea, R.; Moser, H. E. Physicochemical Properties of Antibacterial Compounds: Implications for Drug Discovery. J. Med. Chem. 2008, 51 (10), 2871–2878. https://doi.org/10.1021/jm700967e.

35. Ahlgren, H. G.; Benedetti, A.; Landry, J. S.; Bernier, J.; Matouk, E.; Radzioch, D.; Lands, L. C.; Rousseau, S.; Nguyen, D. Clinical Outcomes Associated with *Staphylococcus aureus* and *Pseudomonas aeruginosa* Airway Infections in Adult Cystic Fibrosis Patients. BMC Pulm. Med. 2015, 15 (1), 67. https://doi.org/10.1186/s12890-015-0062-7.

36. Limoli, D. H.; Yang, J.; Khansaheb, M. K.; Helfman, B.; Peng, L.; Stecenko, A. A.; Goldberg, J. B. *Staphylococcus aureus* and *Pseudomonas aeruginosa* Co-Infection Is Associated with Cystic Fibrosis-Related Diabetes and Poor Clinical Outcomes. Eur. J. Clin. Microbiol. Infect. Dis. 2016, 35 (6), 947– 953. https://doi.org/10.1007/s10096-016-2621-0.

37. Maliniak, M. L.; Stecenko, A. A.; McCarty, N. A. A Longitudinal Analysis of Chronic MRSA and *Pseudomonas aeruginosa* Co-Infection in Cystic Fibrosis: A Single-Center Study. J. Cyst. Fibros. 2016, 15 (3), 350–356. https://doi.org/10.1016/j.jcf.2015.10.014.

38. Hubert, D.; Réglier-Poupet, H.; Sermet-Gaudelus, I.; Ferroni, A.; Bourgeois, M. L.; Burgel, P.-R.; Serreau, R.; Dusser, D.; Poyart, C.; Coste, J. Association between *Staphylococcus aureus* Alone or Combined with *Pseudomonas aeruginosa* and the Clinical Condition of Patients with Cystic Fibrosis. J. Cyst. Fibros. 2013, 12 (5), 497–503. https://doi.org/10.1016/j.jcf.2012.12.003.

39. Ang, M. T. C.; Gumbau-Brisa, R.; Allan, D. S.; McDonald, R.; Ferguson, M. J.; Holbein, B. E.; Bierenstiel, M. DIBI, a 3-Hydroxypyridin-4-One Chelator Iron-Binding Polymer with Enhanced Antimicrobial Activity. MedChemComm 2018, 9 (7), 1206–1212. https://doi.org/10.1039/c8md00192h.

40. Walsh, T. R.; Howe, R. A. The Prevalence and Mechanisms of Vancomycin Resistance in *Staphylococcus aureus*. Annu. Rev. Microbiol. 2002, 56 (1), 657–675. https://doi.org/10.1146/annurev.micro.56.012302.160806.

41. Ropy, A.; Cabot, G.; Sánchez-Diener, I.; Aguilera, C.; Moya, B.; Ayala, J. A.; Oliver, A. Role of *Pseudomonas aeruginosa* Low-Molecular-Mass Penicillin-Binding Proteins in AmpC Expression, β-Lactam Resistance, and Peptidoglycan Structure. Antimicrob. Agents Chemother. 2015, 59 (7), 3925–3934. https://doi.org/10.1128/AAC.05150-14.

42. Torrens, G.; Hernández, S. B.; Ayala, J. A.; Moya, B.; Juan, C.; Cava, F.; Oliver, A. Regulation of AmpC-Driven β-Lactam Resistance in *Pseudomonas aeruginosa*: Different Pathways, Different Signaling. mSystems 2019, 4 (6), e00524–19. https://doi.org/10.1128/mSystems.00524-19.

43. Zamorano, L.; Moyá, B.; Juan, C.; Oliver, A. Differential β-Lactam Resistance Response Driven by AmpD or DacB (PBP4) Inactivation in Genetically Diverse *Pseudomonas aeruginosa* Strains. J. Antimicrob. Chemother. 2010, 65 (7), 1540–1542. https://doi.org/10.1093/jac/dkq142.

44. Barbieri, C. M.; Mack, T. R.; Robinson, V. L.; Miller, M. T.; Stock, A. M. Regulation of Response Regulator Autophosphorylation through Interdomain Contacts. J. Biol. Chem. 2010, 285 (42), 32325–32335. https://doi.org/10.1074/jbc.M110.157164.

45. Boll, J. M.; Hendrixson, D. R. A Specificity Determinant for Phosphorylation in a Response Regulator Prevents in Vivo Cross-Talk and Modification by Acetyl Phosphate. Proc. Natl. Acad. Sci. 2011, 108 (50), 20160–20165. https://doi.org/10.1073/pnas.1113013108.

46. Kurabayashi, K.; Hirakawa, Y.; Tanimoto, K.; Tomita, H.; Hirakawa, H. Role of the CpxAR Two-Component Signal Transduction System in Control of Fosfomycin Resistance and Carbon Substrate Uptake. J. Bacteriol. 2014, 196 (2), 248–256. https://doi.org/10.1128/JB.01151-13.

47. Mahoney, T. F.; Silhavy, T. J. The Cpx Stress Response Confers Resistance to Some, but Not All, Bactericidal Antibiotics. J. Bacteriol. 2013, 195 (9), 1869–1874. https://doi.org/10.1128/JB.02197-12.

48. Masi, M.; Pinet, E.; Pagès, J.-M. Complex Response of the CpxAR Two-Component System to β-Lactams on Antibiotic Resistance and Envelope Homeostasis in Enterobacteriaceae. Antimicrob. Agents Chemother. 2020, 64 (6), e00291–20. https://doi.org/10.1128/AAC.00291-20.

49. Holm, L. Dali Server: Structural Unification of Protein Families. Nucleic Acids Res. 2022, 50 (W1), W210–W215. https://doi.org/10.1093/nar/gkac387.

50. Jumper, J.; Evans, R.; Pritzel, A.; Green, T.; Figurnov, M.; Ronneberger, O.; Tunyasuvunakool, K.; Bates, R.; Žídek, A.; Potapenko, A.; Bridgland, A.; Meyer, C.; Kohl, S. A. A.; Ballard, A. J.; Cowie, A.; Romera-Paredes, B.; Nikolov, S.; Jain, R.; Adler, J.; Back, T.; Petersen, S.; Reiman, D.; Clancy, E.; Zielinski, M.; Steinegger, M.; Pacholska, M.; Berghammer, T.; Bodenstein, S.; Silver, D.; Vinyals, O.; Senior, A. W.; Kavukcuoglu, K.; Kohli, P.; Hassabis, D. Highly Accurate Protein Structure Prediction with AlphaFold. Nature 2021, 596 (7873), 583–589. https://doi.org/10.1038/s41586-021-03819-2.

51. Osawa, T.; Sugiura, N.; Shimada, H.; Hirooka, R.; Tsuji, A.; Shirakawa, T.; Fukuyama, K.; Kimura, M.; Kimata, K.; Kakuta, Y. Crystal Structure of Chondroitin Polymerase from *Escherichia coli* K4. Biochem. Biophys. Res. Commun. 2009, 378 (1), 10–14. https://doi.org/10.1016/j.bbrc.2008.08.121.

52. Kocíncová, D.; Ostler, S. L.; Anderson, E. M.; Lam, J. S. Rhamnosyltransferase Genes MigA and WapR Are Regulated in a Differential Manner To Modulate the Quantities of Core Oligosaccharide Glycoforms Produced by *Pseudomonas aeruginosa*. J. Bacteriol. 2012, 194 (16), 4295–4300. https://doi.org/10.1128/JB.05741-11.

53. Poon, K. K. H.; Westman, E. L.; Vinogradov, E.; Jin, S.; Lam, J. S. Functional Characterization of MigA and WapR: Putative Rhamnosyltransferases Involved in Outer Core Oligosaccharide Biosynthesis of *Pseudomonas aeruginosa*. J. Bacteriol. 2008, 190 (6), 1857–1865. https://doi.org/10.1128/JB.01546-07.

54. Nelson, C. E.; Huang, W.; Zygiel, E. M.; Nolan, E. M.; Kane, M. A.; Oglesby, A. G. The Human Innate Immune Protein Calprotectin Elicits a Multimetal Starvation Response in *Pseudomonas aeruginosa*. Microbiol. Spectr. 2021, 9 (2), e00519–21. https://doi.org/10.1128/Spectrum.00519-21.

55. Kucharczyk, M.; Brzezowska, M.; Maciąg, A.; Lis, T.; Jeżowska-Bojczuk, M. Structural Features of the Cu2+–Vancomycin Complex. J. Inorg. Biochem. 2008, 102 (4), 936–942. https://doi.org/10.1016/j.jinorgbio.2007.12.014.

56. Świątek, M.; Valensin, D.; Migliorini, C.; Gaggelli, E.; Valensin, G.; Jeżowska-Bojczuk, M. Unusual Binding Ability of Vancomycin towards Cu2+ Ions. Dalton Trans. 2005, No. 23, 3808. https://doi.org/10.1039/b508662k.

57. Raivio, T. L.; Silhavy, T. J. Transduction of Envelope Stress in *Escherichia coli* by the Cpx Two-Component System. J. Bacteriol. 1997, 179 (24), 7724–7733. https://doi.org/10.1128/jb.179.24.7724-7733.1997.

58. Hugonnet, J.-E.; Mengin-Lecreulx, D.; Monton, A.; den Blaauwen, T.; Carbonnelle, E.; Veckerlé, C.; Brun, Y. V.; van Nieuwenhze, M.; Bouchier, C.; Tu, K.; Rice, L. B.; Arthur, M. Factors Essential for L,D-Transpeptidase-Mediated Peptidoglycan Cross-Linking and β-Lactam Resistance in *Escherichia coli*. eLife 2016, 5, e19469. https://doi.org/10.7554/eLife.19469.

59. Anderson, E. M.; Shaji Saji, N.; Anderson, A. C.; Brewer, D.; Clarke, A. J.; Khursigara, C. M. *Pseudomonas aeruginosa* Alters Peptidoglycan Composition under Nutrient Conditions Resembling Cystic Fibrosis Lung Infections. mSystems 7 (3), e00156–22. https://doi.org/10.1128/msystems.00156-22.

60. Bielnicki, J.; Devedjiev, Y.; Derewenda, U.; Dauter, Z.; Joachimiak, A.; Derewenda, Z. S. B. Subtilis YkuD Protein at 2.0 Å Resolution: Insights into the Structure and Function of a Novel, Ubiquitous Family of Bacterial Enzymes. Proteins 2006, 62 (1), 144–151. https://doi.org/10.1002/prot.20702.

61. Dbeibo, L.; van Rensburg, J. J.; Smith, S. N.; Fortney, K. R.; Gangaiah, D.; Gao, H.; Marzoa, J.; Liu, Y.; Mobley, H. L. T.; Spinola, S. M. Evaluation of CpxRA as a Therapeutic Target for Uropathogenic *Escherichia coli* Infections. Infect. Immun. 2018, 86 (3), e00798–17. https://doi.org/10.1128/IAI.00798-17.

62. DeLeon, S.; Clinton, A.; Fowler, H.; Everett, J.; Horswill, A. R.; Rumbaugh, K. P. Synergistic Interactions of *Pseudomonas aeruginosa* and *Staphylococcus aureus* in an In Vitro Wound Model. Infect. Immun. 2014, 82 (11), 4718–4728. https://doi.org/10.1128/IAI.02198-14.

63. Murdoch, C. C.; Skaar, E. P. Nutritional Immunity: The Battle for Nutrient Metals at the Host– Pathogen Interface. Nat. Rev. Microbiol. 2022, 20 (11), 657–670. https://doi.org/10.1038/s41579-022-00745-6.

64. P. Skaar, E.; Raffatellu, M. Metals in Infectious Diseases and Nutritional Immunity. Metallomics 2015, 7 (6), 926–928. https://doi.org/10.1039/C5MT90021B.

65. Stokes, J. M.; French, S.; Ovchinnikova, O. G.; Bouwman, C.; Whitfield, C.; Brown, E. D. Cold Stress Makes *Escherichia coli* Susceptible to Glycopeptide Antibiotics by Altering Outer Membrane Integrity. Cell Chem. Biol. 2016, 23 (2), 267–277. https://doi.org/10.1016/j.chembiol.2015.12.011.

66. Belanger, C. R.; Lee, A. H.-Y.; Pletzer, D.; Dhillon, B. K.; Falsafi, R.; Hancock, R. E. W. Identification of Novel Targets of Azithromycin Activity against *Pseudomonas aeruginosa* Grown in Physiologically Relevant Media. Proc. Natl. Acad. Sci. U. S. A. 2020, 117 (52), 33519–33529. https://doi.org/10.1073/pnas.2007626117.

67. Chibani-Chennoufi, S.; Bruttin, A.; Dillmann, M.-L.; Brüssow, H. Phage-Host Interaction: An Ecological Perspective. J. Bacteriol. 2004, 186 (12), 3677–3686. https://doi.org/10.1128/JB.186.12.3677-3686.2004.

68. Loh, B.; Grant, C.; Hancock, R. E. Use of the Fluorescent Probe 1-N-Phenylnaphthylamine to Study the Interactions of Aminoglycoside Antibiotics with the Outer Membrane of *Pseudomonas aeruginosa*. Antimicrob. Agents Chemother. 1984, 26 (4), 546–551. https://doi.org/10.1128/AAC.26.4.546.

69. Liberati, N. T.; Urbach, J. M.; Miyata, S.; Lee, D. G.; Drenkard, E.; Wu, G.; Villanueva, J.; Wei, T.; Ausubel, F. M. An Ordered, Nonredundant Library of *Pseudomonas aeruginosa* Strain PA14 Transposon Insertion Mutants. Proc. Natl. Acad. Sci. U. S. A. 2006, 103 (8), 2833–2838. https://doi.org/10.1073/pnas.0511100103.

70. Feng, M.; Mei, J.; Hu, S.; Janney, S.; Carruthers, J.; Holbein, B.; Huber, A.; Kidby, D. Selective Removal of Iron from Grape Juice Using an Iron(III) Chelating Resin. Sep. Purif. Technol. 1997, 11 (2), 127–135. https://doi.org/10.1016/S1383-5866(97)00010-5.

71. Mangat, C. S.; Bharat, A.; Gehrke, S. S.; Brown, E. D. Rank Ordering Plate Data Facilitates Data Visualization and Normalization in High-Throughput Screening. J. Biomol. Screen. 2014, 19 (9), 1314–1320. https://doi.org/10.1177/1087057114534298.

72. Deatherage, D. E.; Barrick, J. E. Identification of Mutations in Laboratory Evolved Microbes from Next-Generation Sequencing Data Using Breseq. Methods Mol. Biol. Clifton NJ 2014, 1151, 165–188. https://doi.org/10.1007/978-1-4939-0554-6_12.

73. Marolda, C. L.; Lahiry, P.; Vinés, E.; Saldías, S.; Valvano, M. A. Micromethods for the Characterization of Lipid A-Core and O-Antigen Lipopolysaccharide. In Glycobiology Protocols; Brockhausen, I., Ed.; Methods in Molecular Biology; Humana Press: Totowa, NJ, 2007; pp 237–252. https://doi.org/10.1385/1-59745-167-3:237.

74. Kocíncová, D.; Hao, Y.; Vinogradov, E.; Lam, J. S. Evidence That WapB Is a 1,2-Glucosyltransferase of *Pseudomonas aeruginosa* Involved in Lipopolysaccharide Outer Core Biosynthesis. J. Bacteriol. 2011, 193 (11), 2708–2716. https://doi.org/10.1128/JB.00032-11.

75. Harvey, H.; Bondy-Demony, J.; Marquis, H.; Sztanko, K. M.; Davidson, A. R.; Burrows, L. L. *Pseudomonas aeruginosa* defends against phages through type IV pilus glycosylation. Nat. Microbiol. 2018. 3, 47–52. https://doi.org/10.1038/s41564-017-0061-y.

76. Goddard, T. D.; Huang, C. C.; Meng, E. C.; Pettersen, E. F.; Couch, G. S.; Morris, J. H.; Ferrin, T. E. UCSF ChimeraX: Meeting Modern Challenges in Visualization and Analysis. Protein Sci. 2018, 27 (1), 14–25. https://doi.org/10.1002/pro.3235.

