## Supplementary Figures S1-S7 and Tables S1-S4 for "Nutrient Limitation Sensitizes *Pseudomonas aeruginosa* to Vancomycin"

**Supplementary Data for: Nutrient limitation sensitizes *Pseudomonas aeruginosa* to vancomycin**

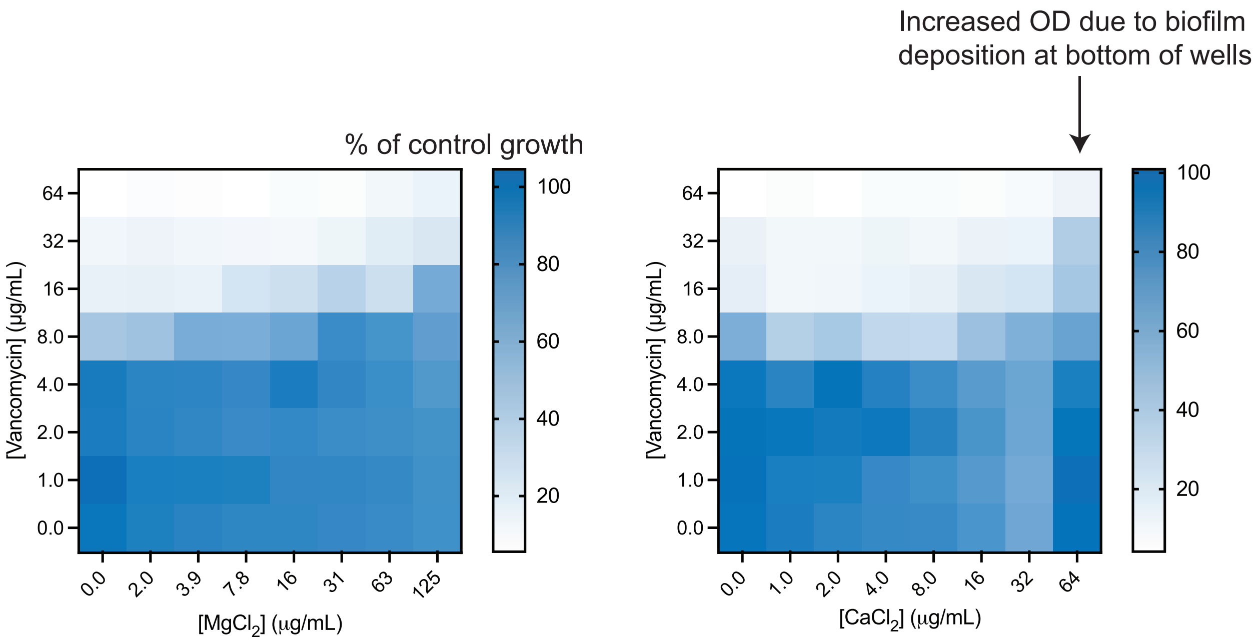


**Supplementary Figure S1.** Checkerboard assays of vancomycin with $MgCl_2$ and $CaCl_2$ against PA14. Results are averaged from three independent biological replicates.

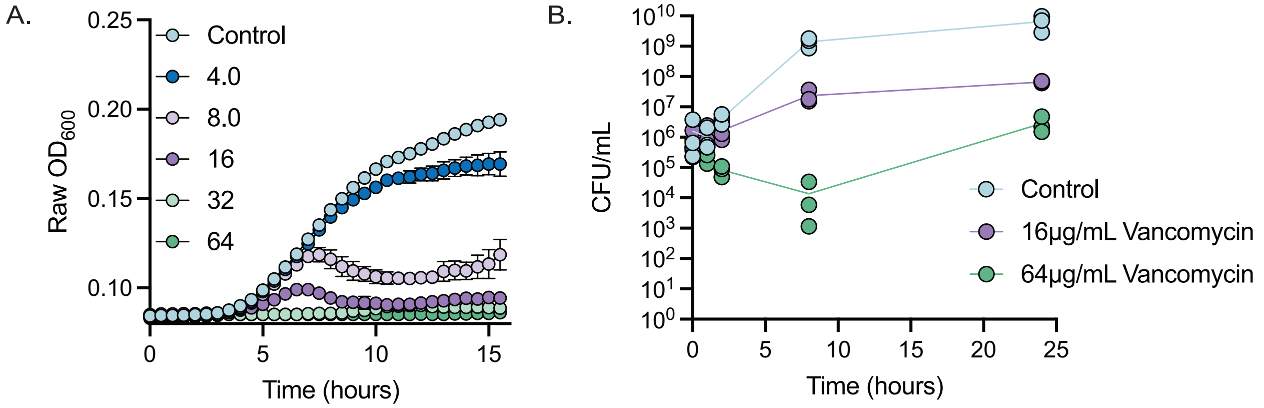


**Supplementary Figure S2. A.** Growth curves with PA14 treated with increasing concentrations of vancomycin (µg/mL) and **B.** time-kill kinetics of vancomycin at 1X MIC (16µg/mL) and 4X MIC (64µg/mL) in 10:90. All results are averaged from three independent biological replicates. Individual biological replicates are shown for the time-kill kinetics. Cells were grown in 10:90 for all assays.

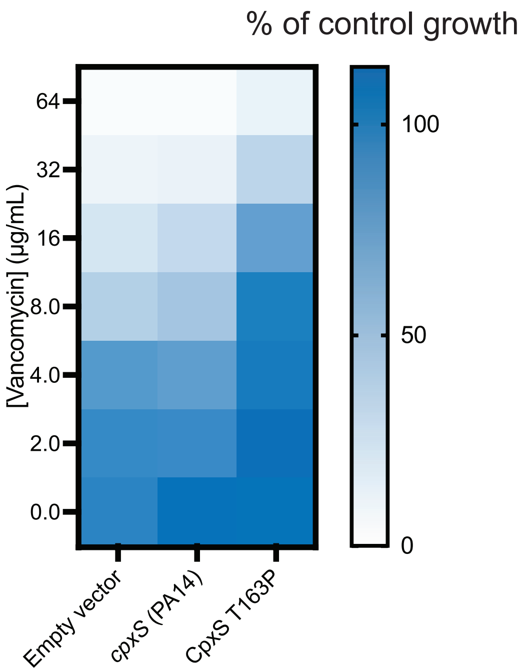


**Supplementary Figure S3.** Vancomycin MIC assays with PA14 *cpxS*::Mar2xT7 with empty vector, expressing WT PA14 *cpxS,* and CpxS T163P in 10:90 + 0.25% arabinose. Results are averaged from three independent biological replicates.

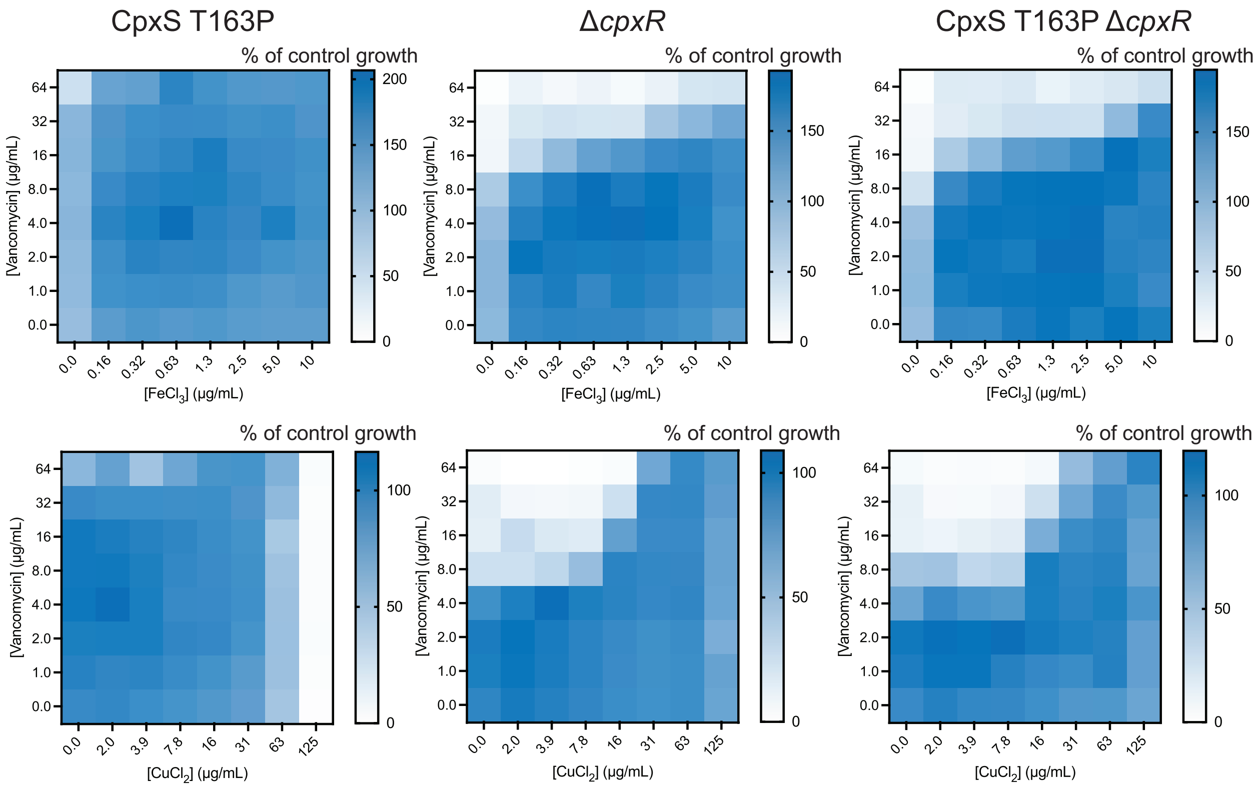


**Supplementary Figure S4.** Checkerboard assays of vancomycin with $FeCl_3$ and $CuCl_2$ against Δ*cpxR,* CpxS T163P, and CpxS T163P Δ*cpxR*. Results are averaged from three independent biological replicates.

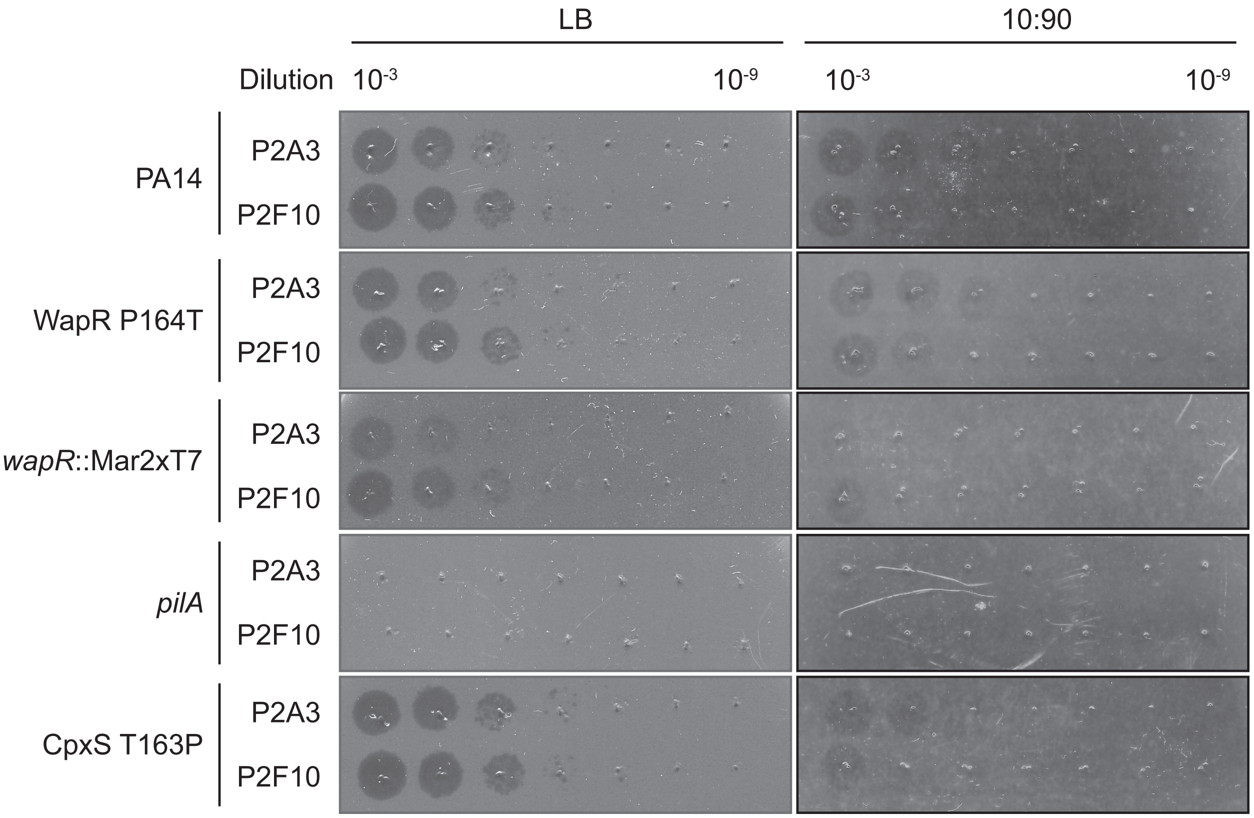


**Supplementary Figure S5.** WapR P164T mutant is not sensitized to pilus-specific phages in LB or 10:90.

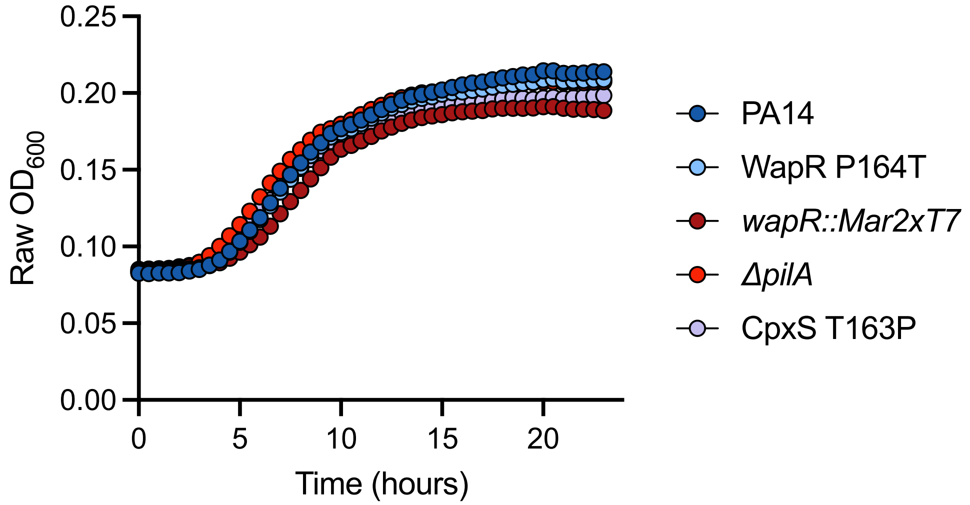


**Supplementary Figure S6.** Growth curves of PA14, WapR P164T, *wapR*::Mar2xT7, Δ*pilA*, and CpxS T163P in 10:90. Results are averaged from three independent biological replicates.

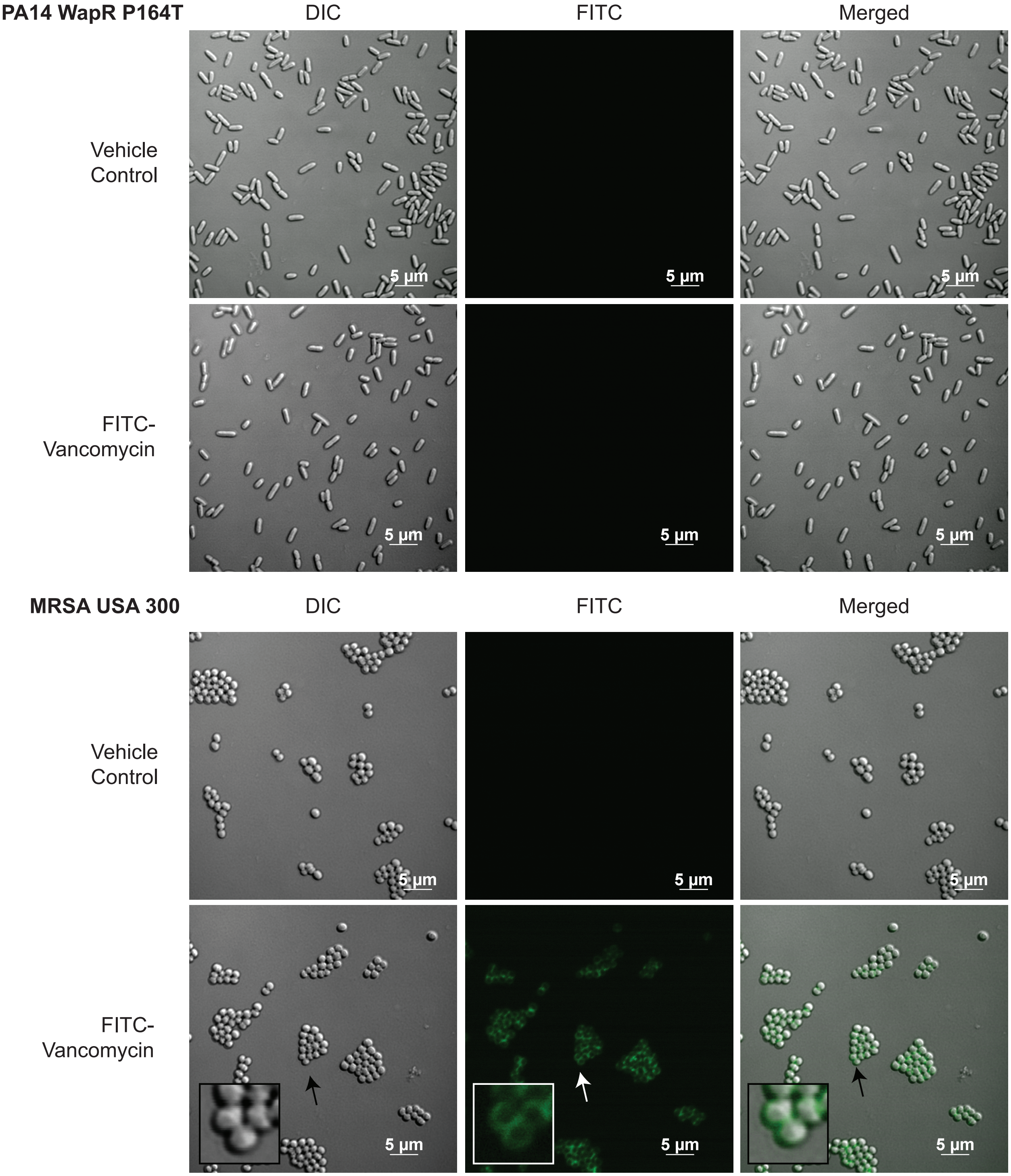


**Supplementary Figure S7.** WapR P164T is not labeled by FITC-vancomycin. Labeling of MRSA USA 300 is shown as a positive control. Representative images are shown. Scale bar is indicated.

**Supplementary Table S1**: List of Compounds in the Initial Screen

| Compound | Activity against PA14 | Molecular Weight (g/mol) |
|---|---|---|
| A54556A | N | 718.8 |
| Actagardin | N | 1754 |
| Actinomycin D | Y | 1255.4 |
| Amphomycin | N | 1290.4 |
| Aspartocin D | N | 1276.4 |
| Azithromycin | Y | 749 |
| Boromycin | N | 879.9 |
| Capreomycin | N | 668.7 |
| Echinomycin | Y | 1101.3 |
| Feglymycin | N | 1900.9 |
| Fidaxomicin | N | 1058.1 |
| Gramicidin | N | 1882.3 |
| Luzopeptin A | N | 1343.3 |
| Nosiheptide | Y | 1222.4 |
| Quinaldopeptin | Y | 1243.3 |
| Quinupristin | N | 1022.2 |
| Sandramycin | Y | 1221.3 |
| Siamycin | N | 2163.5 |
| Streptogramin B | N | 867 |
| Telomycin | N | 1272.3 |
| Thiocoraline | N | 1157.4 |
| Tunicamycin | N | 816.9 |
| Virginiamycin | N | 1349.5 |

Y: has antimicrobial activity against *P. aeruginosa* PA14 in the screen (highlighted blue)
N: lacks antimicrobial activity against *P. aeruginosa* PA14 in the screen (highlighted red)

**Supplementary Table S2**. Strains used in this study

| Strain | Source |
|---|---|
| PA14 | 1 |
| PA14 Δ*cpxR* | This study |
| PA14 CpxS T163P | This study |
| PA14 CpxS T163P Δ*cpxR* | This study |
| PA14 Δ*cpxR* pBADGr | This study |
| PA14 Δ*cpxR* pBADGr-*cpxR* | This study |
| PA14 CpxS T163P Δ*cpxR* pBADGr | This study |
| PA14 CpxS T163P Δ*cpxR* pBADGr-*cpxR* | This study |
| PA14 WapR P164T | This study |
| PA14 *wapR*::Mar2xT7 | 1 |

| | |
|---|---|
| PA14 *cpxS*::Mar2xT7 | 1 |
| PA14 *cpxS*::Mar2xT7 pHERD20T | This study |
| PA14 *cpxS*::Mar2xT7 pHERD20T-*cpxS*$_{PA14}$ | This study |
| PA14 *cpxS*::Mar2xT7 pHERD20T-*cpxS*$_{PA14}$ T163P | This study |
| PA14 Δ*pilA* | This study |
| PA14 Δ*dacB* | This study |
| PA14 Δ*dacC* | This study |
| PA14 Δ*dacB* Δ*dacC* | This study |
| PA14 P$_{x}$-*gfp* | 2 |
| PA14 P$_{mexA}$-*gfp* | This study |
| PA14 CpxS T163P P$_{x}$-*gfp* | This study |
| PA14 CpxS T163P P$_{mexA}$-*gfp* | This study |
| *P. aeruginosa* C0275 | 3 |
| *P. aeruginosa* C0098 | 3 |
| MRSA USA300 | Brown Lab |

**Supplementary Table S3.** Primers used in this study. Restriction sites are bolded and underlined.

| Primer | Forward 5' → 3' | Reverse 5' → 3' |
|---|---|---|
| *cpxR* deletion upstream | CCT**GGATCC**GTGGTCTTCACCTATCCGAACTTCTGG | TCAGTGGCTGTAGTAGTAGCCGCGGCAGAGCTCCCGGTCATCG |
| *cpxR* deletion downstream | CGATGACCGGGAGCTCTGCCGCGGCTACTACTACAGCCACTGA | CCT**TCTAGA**TCCTGTTGAATCGCATGTCGGG |
| CpxS T163P chromosomal mutation upstream | CAT**GAATTC**GGCCCGACATGCGATTCAACAG | TGGTGGTGCTGCCTCTGTTCA |
| CpxS T163P chromosomal mutation downstream | TGAACAGAGGCAGCACCACCA | CCC**AAGCTT**TTATTCGGTCGCCTTGCGCC |
| *cpxR* complementation | CGT**GAATTC**CGACAGCGACGCGAATGGATC | GCT**AAGCTT**GCTTGGGTAAAGACCTTGTGTCGG |
| *dacB* chromosomal deletion upstream | CGT**TCTAGA**AGCAATTGGCCTGGTTATCTGCATGA | TTCGTACTGCGCCTTATTTCCGCGTAATGAAGAACCCTTGAAGACGGAGGGTAG |
| *dacB* chromosomal deletion downstream | CTACCCTCCGTCTTCAAGGGTTCTTCATTACGCGGAAATAAGGCGCAGTACGAA | CAC**AAGCTT**AGGAGCGGGTGGTGGTCAAG |
| *dacC* chromosomal deletion upstream | GCT**TCTAGA**GACTATTCCCGTCCGCATCGC | GGTCGGTCAGTTGAACAAACCGTAGAAATTCCCTGCGTGAGTAGGCGC |
| *dacC* chromosomal deletion downstream | GCGCCTACTCACGCAGGGAATTTCTACGGTTTGTTCAACTGACCGACC | CAG**AAGCTT**TGCTCGATCCGGCTGACCA |
| P$_{mexA}$-*gfp* | CCA**TCTAGA**GCTCGCGGATCTTCCGGGT | CAAT**GCATGC**TCAGCGAAAGCGGCCGATGCA |
| *cpxS* amplification | CAT**GAATTC**GGCCCGACATGCGATTCAACAG | CCC**AAGCTT**TTATTCGGTCGCCTTGCGCC |
| *pilA* chromosomal deletion upstream | TCGA**GGATCC**GCATCACGATCTTCTC | TAC**CTGCAG**TCGCAACCACGATCATCAG |
| *pilA* chromosomal deletion downstream | AC**CTGCAG**ATATGCCTGCCCTGACTGCA | CTGG**AAGCTT**CCGGCGGAATCAACG |

**Supplementary Table S4**: Compounds used in this study

| **Vendor/Source** | **Compound** |
|---|---|
| Bioshop | Kanamycin |
| Bioshop | Gentamicin |
| Bioshop | Arabinose |
| AK Scientific | Carbenicillin |
| AK Scientific | Vancomycin |
| AK Scientific | Imipenem |
| AK Scientific | Meropenem |
| AK Scientific | Tobramycin |
| AK Scientific | Azithromycin |
| AK Scientific | $CuCl_2$ |
| Sigma | FITC-vancomycin |
| Sigma | Polymyxin B |
| Sigma | Cefmetazole |
| Sigma | Cefoperazone |
| Sigma | Aztreonam |
| Sigma | Ceftriaxone |
| Sigma | Piepracillin |
| Sigma | Cefuroxime |
| Sigma | Cefsulodin |
| Sigma | A22 |
| Sigma | Fosfomycin |
| Sigma | Doxycycline |
| Sigma | Ciprofloxacin |
| Sigma | Novobiocin |
| Cayman Chemicals | D-cycloserine |
| Cayman Chemicals | Dalbavancin |
| Cayman Chemicals | Oritavancin |
| Cayman Chemicals | Teicoplanin |
| Cayman Chemicals | CCCP |
| Cayman Chemicals | Nosiheptide |
| Cayman Chemicals | Actinomycin D |
| Cayman Chemicals | Echinomycin |
| Cayman Chemicals | Virginiamycin |
| Cayman Chemicals | Amphomycin |
| Cayman Chemicals | Capreomycin |
| Cayman Chemicals | Fidaxomicin |
| Cayman Chemicals | A54556A |
| Cayman Chemicals | Streptogramin B |
| Cayman Chemicals | Quinupristin |
| Cayman Chemicals | Gramicidin |
| Cayman Chemicals | Tunicamycin |
| Cayman Chemicals | Telomycin |
| Cayman Chemicals | Boromycin |

| | |
|---|---|
| Cayman Chemicals | Siamycin |
| Cayman Chemicals | Astagardin |
| Cayman Chemicals | Thiocoraline |
| Cayman Chemicals | Luzopeptin A |
| Cayman Chemicals | Feglymicin |
| Cayman Chemicals | Sandramycin |
| Cayman Chemicals | Quinaldopeptin |
| Cayman Chemicals | Aspartocin D |
| Fisher Scientific | NPN |
| Fisher Scientific | $FeCl_3$ |
| DIBI | Dr. Bruce Holbein (Fe Pharmaceuticals) |
| FEC-1 (insoluble DIBI analog to remove iron from media) | Dr. Bruce Holbein (Fe Pharmaceuticals) |
| MedChemExpress | Cefiderocol |

**SUPPLEMENTARY REFERENCES**


1. Liberati, N. T.; Urbach, J. M.; Miyata, S.; Lee, D. G.; Drenkard, E.; Wu, G.; Villanueva, J.; Wei, T.; Ausubel, F. M. An Ordered, Nonredundant Library of *Pseudomonas aeruginosa* Strain PA14 Transposon Insertion Mutants. *Proc. Natl. Acad. Sci. U. S. A.* **2006**, *103* (8), 2833–2838.
2. Chan, D. C. K.; Josts, I.; Koteva, K.; Wright, G. D.; Tidow, H.; Burrows, L. L. A Single Extracellular Loop of FoxA Controls Ligand Specificity, Uptake, and Signaling in *Pseudomonas aeruginosa*. bioRxiv November 18, 2022, p 2022.11.18.517105.
3. Ranieri, M. R. M.; Chan, D. C. K.; Yaeger, L. N.; Rudolph, M.; Karabelas-Pittman, S.; Abdo, H.; Chee, J.; Harvey, H.; Nguyen, U.; Burrows, L. L. Thiostrepton Hijacks Pyoverdine Receptors To Inhibit Growth of *Pseudomonas aeruginosa*. *Antimicrob. Agents Chemother.* **2019**, *63* (9), e00472-19.
